# CCZ1 is a modulator of TPC2 activity and melanoma cell migration

**DOI:** 10.64898/2026.04.22.718428

**Authors:** Zhuo Yang, Colin Feldmann, Lina Ouologuem, Alice C. Lin, Stefanie Fenske, Stylianos Michalakis, Karin Bartel, Michael Schänzler, Christian Grimm, Cheng-Chang Chen, Christian Wahl-Schott, Martin Biel

**Author notes:** These authors contributed equally (Zhuo Yang, Colin Feldmann). Corresponding authors: Martin Biel; Christian Wahl-Schott, Cheng-Chang Chen.

## Abstract

The small GTPase RAB7a is a key regulator of melanoma progression by enhancing the activity of the endolysosomal two-pore cation channel TPC2. In this study, we demonstrate that CCZ1—a core component of the RAB7a guanine nucleotide exchange factor (GEF) complex—is essential for mediating this RAB7a-dependent enhancement of TPC2. Unexpectedly, we find that constitutively active (GTP-locked) RAB7a fails to bind and regulate TPC2 in the absence of CCZ1, indicating that CCZ1 contributes to the RAB7a–TPC2 interaction through mechanisms beyond its GEF activity. Furthermore, the CCZ1 facilitated GTPase-activating function on RAB5 is dispensable for modulating TPC2. Notably, in the absence of CCZ1, TPC2 exhibits increased affinity for its agonist, PI(3,5)P₂, along with markedly upregulated channel activity. In melanoma cell lines, this upregulation enhances migratory capacity. Our findings identify CCZ1 as a functional inhibitor of TPC2 and highlight its critical role in regulating cancer cell migration.

## Introduction

Two-pore channels (TPCs) are Ca²⁺- and Na⁺-permeable cation channels that are widely expressed throughout the body and are localized to endolysosomal organelles (*1–4*). In both mice and humans, two TPC isoforms exist: TPC1 and TPC2 (*5–7*). TPCs have been implicated in the regulation of various endolysosomal trafficking pathways (*1–4*) and are associated with several pathological processes, including cancer cell migration (*8–11*), neoangiogenesis (*12, 13*), as well as metabolic (*14, 15*) and infectious diseases (*16, 17*). Although both TPCs are expressed intracellularly, TPC1 is predominantly localized to early endosomes (*18*), whereas TPC2 is mainly found in late endosomes and lysosomes (*5*). TPC2 can be activated either directly by phosphatidylinositol 3,5-bisphosphate [PI(3,5)P₂] or indirectly by nicotinic acid adenine dinucleotide phosphate (NAADP), which requires accessory binding proteins including JPT2 (*19, 20*), Lsm12 (*21*), or ASPDH (*22*) to exert its agonistic effect.

The signaling pathways that govern TPC activation remain incompletely understood. Recent studies have identified the small GTPase RAB7a as part of the TPC2 interactome (*23*). Notably, overexpression of RAB7a enhances TPC2-mediated currents, indicating that RAB7a functions as a positive modulator of TPC2 activity (*24*). Functionally, this RAB7a-dependent regulation of TPC2 has been linked to the migration of melanoma cells in both in vitro and in vivo models (*24*).

Small GTPases are activated by the exchange of GDP for GTP, a reaction catalyzed by guanine nucleotide exchange factors (GEFs). The trimeric CCZ1–MON1a/b–RMC1 complex is the only known GEF for RAB7a (*25*). Structural and functional studies have shown that CCZ1 and one of the two MON1 isoforms, MON1a and MON1b, (collectively designated MON1a/b in the following) form the catalytic interface for RAB7a, whereas RMC1 (also known as C18orf8 or Bulli in Drosophila) stabilizes the CCZ1–MON1a/b core but is dispensable for the GDP–GTP exchange reaction itself (*26–29*). Consistent with this supportive role, RMC1 is found only in metazoans, while the CCZ1–MON1a/b core is conserved across all eukaryotes, including unicellular organisms (*30*). Given the central role of CCZ1–MON1a/b in RAB7a activation, we investigated whether this complex is also required for RAB7a-dependent modulation of TPC2. To this end, we disrupted the CCZ1-MON1a/b-RMC1 complex by deleting the central CCZ1 subunit in HEK293 cell lines. In the absence of CCZ1, RAB7a no longer bound to TPC2, nor was it able to activate TPC2-mediated currents. Surprisingly, this loss of modulation was also observed using a constitutively active RAB7a mutant RAB7a^Q67L^ that mimics the GTP-bound form and bypasses the need for CCZ1–MON1a/b-mediated nucleotide exchange (*31*). Taken together, these findings indicate that RAB7a-dependent modulation of TPC2 strictly requires the presence of CCZ1, via a mechanism that is independent of its canonical GEF activity. Unexpectedly, we also observed that in the absence of CCZ1, TPC2 currents were strongly enhanced, reaching levels comparable to those of wild-type (WT) cells upon RAB7a activation. Consistent with this constitutive activation, a CRISPR-Cas9–engineered CCZ1 knockout (KO) melanoma cell line (SK-MEL-5) displayed elevated TPC2 activity and increased migratory behavior.

These results uncover a previously unrecognized functional link between CCZ1 and TPC2, which may play a significant role in cancer progression and other (patho)physiological processes involving TPC2.

## Results

### CCZ1 affects number and size of endolysosomal vesicles

We employed HEK293 cells to investigate the potential role of CCZ1 in modulating TPC2. As these cells lack endogenous TPC2 expression, they are frequently used to characterize the channel following heterologous overexpression (*32*). In contrast, CCZ1 is expressed endogenously in HEK293 cells (Fig. 1A). To evaluate the contribution of CCZ1 to TPC2 modulation, we used CRISPR-Cas9 to disrupt the CCZ1 coding region, effectively abolishing its endogenous expression (Fig. 1A, Fig. S1A, B). Confocal super-resolution (AiryScan) imaging of TPC2-mVenus–overexpressing cells revealed that CCZ1 KO cells contained fewer TPC2-positive endolysosomal organelles per cell, with increased perimeter compared to WT (Fig. 1B, C). This enlargement was confirmed by transmission electron microscopy in non-overexpressing cells, excluding TPC2 overexpression as the cause for the endolysosomal enlargement (Fig. 1D, E). Immunofluorescence staining of WT and CCZ1 KO cells using the early endosomal marker EEA1 and the late endosomal marker LAMP1 revealed that CCZ1 KO cells retain two spatially distinct endosomal populations with minimal overlap (Fig. 1F, G). To further characterize the identity of the enlarged vesicles, cells were co-transfected with TPC2-mTq2 and the endolysosomal marker LAMP1-mVenus. These experiments showed a high degree of fluorescence signal colocalization, indicating that the most prominently enlarged vesicles correspond to late endosomes and corroborating the results shown in Fig. 1F and G (Fig. 1H, I). Importantly, this observation was independent of apilimod treatment (Fig. S1D, E).

**Figure 1:**
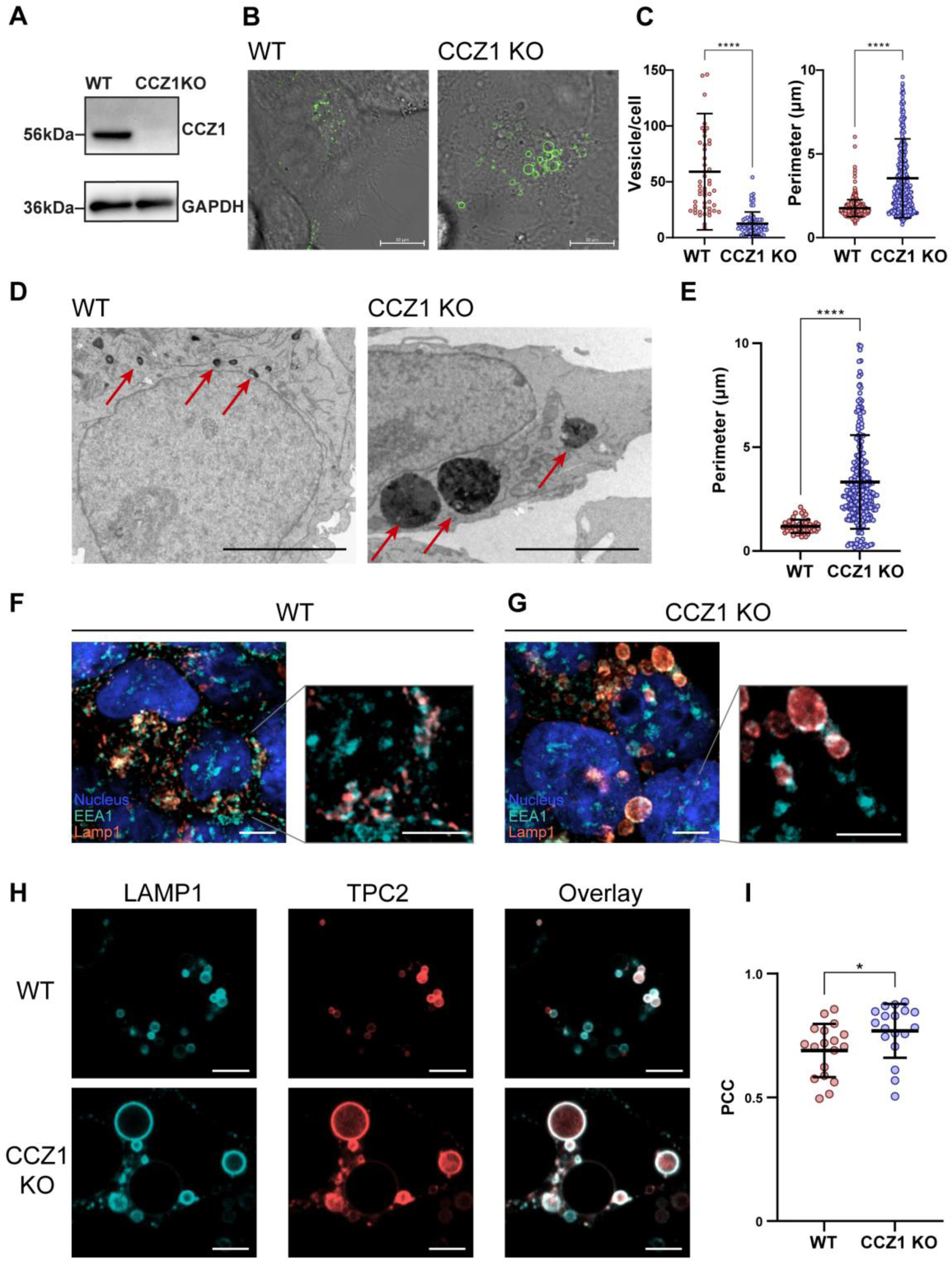
Decreased number and increased perimeter of endosomal vesicles in CCZ1 KO cells. (**A**) Western Blot validation of CCZ1 knockout. (**B**) Live cell confocal super-resolution (AiryScan) images of HEK293 WT and CCZ1 KO cells expressing TPC2-mVenus (green). (**C**) Quantification of the perimeter of TPC2-mVenus–positive vesicles in WT and CCZ1 KO cells (left diagram) and vesicles/cell (right diagram). Each dot represents a TPC2-positive vesicle; mean ± SD is indicated. Statistical significance was assessed using Mann-Whitney test, ****p<0.0001. (**D**) Electron microscopy (TEM) of native WT and CCZ1 KO HEK293 cells. Red arrows indicate potential vesicles (dark, electron-rich areas). Scalebar: 5 µm (**E**) Quantification of vesicle perimeter from objects as indicated by the arrows in D (WT: n=55 CCZ1 KO: n=275). Statistical analysis: Mann-Whitney test, ****p<0.0001. (**F, G**) Immunofluorescence staining of native HEK293 WT (F) and CCZ1 KO cells (G) with early endosomal marker EEA1 (cyan), late endosomal marker LAMP1 (red) and Hoechst (blue). Scale bar: 5 µm. (**H**) Representative confocal images of WT (top) and CCZ1 KO cells (bottom) co-expressing LAMP1-mVenus (cyan) and TPC2-mTq2 (red) treated overnight with apilimod (1 µM). Scalebar: 5 µm (**I**) Quantification of Pearson’s correlation coefficient between TPC2 and LAMP1 signals in multiple cells (n=18) as shown in F. Each dot represents an individual cell; mean ± SD is indicated. Statistical significance was assessed using an unpaired two-tailed t-test, *p<0.05.

### CCZ1 is required for interaction of TPC2 with RAB7a

We next analyzed which effect the deletion of CCZ1 has on the interaction between TPC2 and RAB7a. To this end, prior to analysis, the cells were treated overnight with 1 μM apilimod, a specific PIKfyve inhibitor to enlarge late endosomes and lysosomes (*32*). In agreement with our earlier findings, organellar RAB7a and TPC2 expression pattern strongly overlapped and resulted in high FRET efficiencies using a lifetime-based FRET approach (Fig. 2A upper row, Fig. 2B). These findings were validated further using the intensity based Two-Hybrid FRET method (*33*) (Fig. S2A). In contrast, there was almost no co-localization in CCZ1 KO cells (Fig. 2A, lower row, Fig. 2B and Fig. S2C) and FRET signals were abolished. To assess the specificity of the RAB7a–TPC2 interaction and exclude crowding–driven false–positive FRET signals, we analyzed RAB7a and LAMP1 as a control FRET pair and detected no measurable FRET signal (Fig. S3A, B)

**Figure 2:**
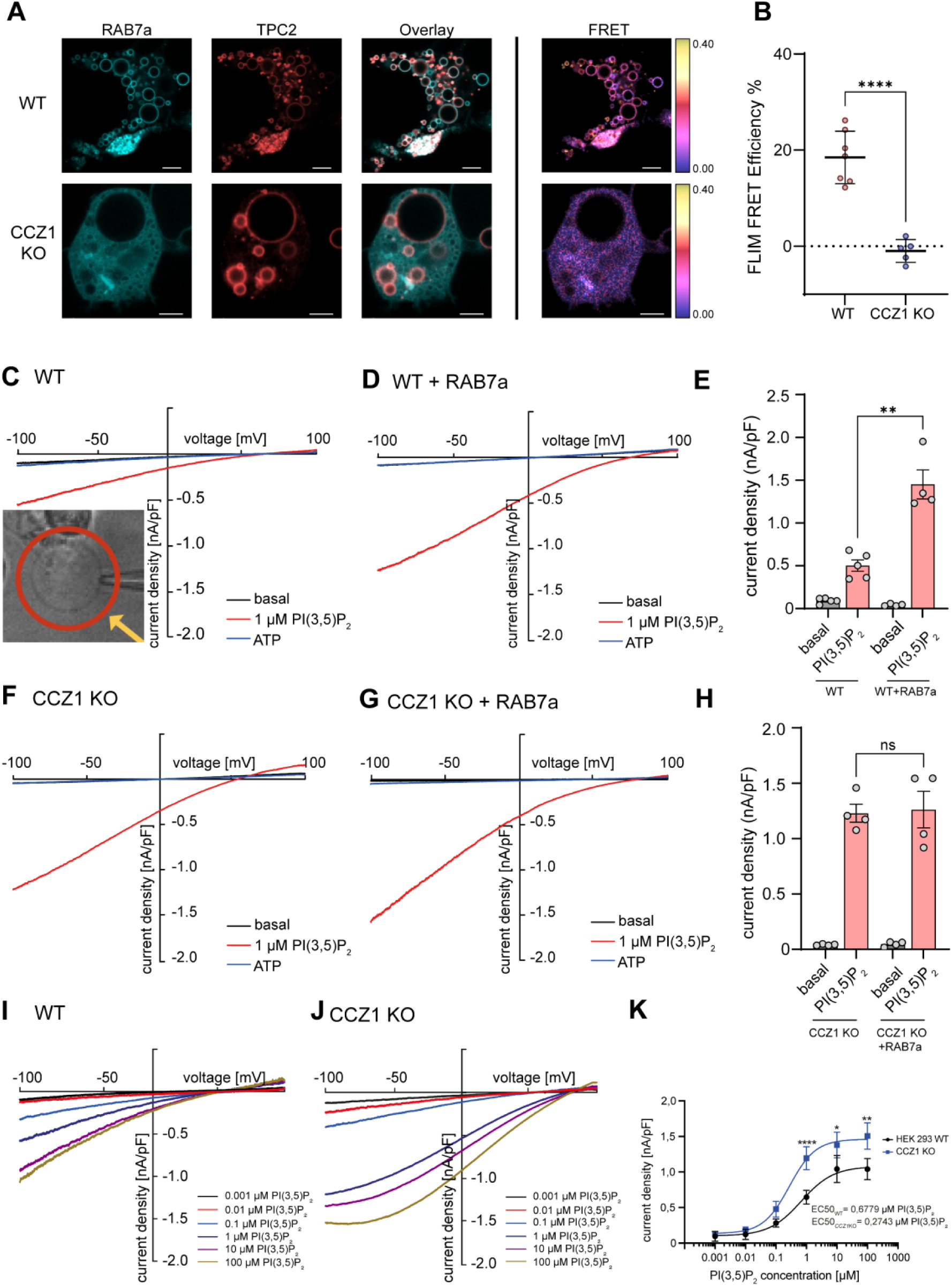
CCZ1 KO disrupts the RAB7a–TPC2 interaction and impairs RAB7a-dependent TPC2 channel activity. (**A**) HEK293 WT (top row) and CCZ1 KO cells co-expressing mTq2-RAB7a (cyan) and TPC2-mVenus (red), treated with apilimod (1 µM), with corresponding FLIM-FRET efficiency maps shown on the right. FRET efficiencies (0–40%) are color-coded according to the color scale. Scale bars: 5 µm. (**B**) FRET efficiencies measured in representative regions of interest (FRET-positive vesicles in WT cells and representative mTq2-RAB7a-positive areas in CCZ1 KO cells) corresponding to images in A (n=7 WT, n=5 CCZ1 KO). Error bars: ±SD. Statistical significance: Unpaired two tailed t-test, ****p<0.0001. (**C-D**, **F-G**) Whole-endolysosomal patch-clamp recordings of overexpressed TPC2 currents in HEK293 WT and CCZ1 KO cells, showing basal (black), ATP (blue), and PI(3,5)P₂ (red)-stimulated current densities. Cells were pretreated with apilimod (1 µM) overnight. An isolated endosome in bath solution during patch-clamp is shown in the left bottom of panel C. (**C-D**) Representative current-voltage (I-V) relationships in WT cells in the absence (C) and presence (D) of RAB7a co-expression. (**E**) Quantification of PI(3,5)P_2_-mediated TPC2 current densities at -100 mV from endolysosomal recordings in (C**-**D), n=4 for each condition. Data are presented as mean ± SEM, statistical significance: unpaired two-tailed t-test, **p<0.01 (p=0.0067). (**F-G**) Representative current-voltage (I-V) relationships in CCZ1 KO cells in the absence (F) and presence (G) of RAB7 co-expression. (**H**) Quantification of PI(3,5)P_2_-mediated TPC2 current densities at -100 mV from endolysosomal recordings in (F, G). Data are presented as mean ± SEM. Each dot on the bar graph represents a single current density value measured from one endolysosome (EL). Statistical significance: unpaired two tailed t-test. ns, not significant. (**I-J**) Whole-endolysosome patch-clamp recordings of overexpressed TPC2 currents in WT (I) and CCZ1 KO (J) HEK293 cells in response to varying concentrations of PI(3,5)P₂ (0.001 μM, 0.01 μM, 0.1 μM, 1 μM, 10 μM, 100 μM). (**K**) Dose-response curve of overexpressed TPC2 comparing the current densities at -100mV induced by increasing concentrations of PI(3,5)P₂ in WT (black) and CCZ1 KO (blue) cells. At least 4 recordings were obtained for each concentration. Data are presented as mean ± SEM, statistical significance was determined using unpaired t-test (two-tailed), 1 μM ****p<0.0001, 10 μM *p=<0.1 (p=0.0163), 100 μM **p=<0.01 (p=0.0057).

We next examined whether CCZ1 functionally interferes with RAB7a effects on TPC2 currents by employing endolysosomal patch-clamp experiments (Fig. 2 C–H). These experiments demonstrated that robust TPC2-mediated cation currents could be activated by the agonist PI(3,5)P₂ in both WT and CCZ1 KO cells. In both cases, the well-known TPC inhibitor ATP (*34*) effectively abolished the currents. Consistent with previous findings, RAB7a significantly enhanced TPC2-mediated currents in WT cells (Fig 2 C–E). In contrast, RAB7a had no stimulatory effect on TPC2 currents in CCZ1 KO cells (Fig. 2 F–H).

Strikingly, TPC2 current densities were markedly increased in CCZ1 KO cells compared to WT cells and were comparable to those in WT cells co-expressing TPC2 and RAB7a. The normalized TPC2 current densities were 0.50 ± 0.067 nA/pF (n = 5) in WT, 1.45 ± 0.17 nA/pF (n = 5) in WT + RAB7a, 1.23 ± 0.08 nA/pF (n = 5) in CCZ1 KO, and 1.26 ± 0.16 nA/pF (n = 4) in CCZ1 KO + RAB7a. To determine whether the increased current density in CCZ1 KO cells reflects a general effect on endolysosomal cation channels, we analyzed endogenous TRPML1 currents. TRPML1 belongs to the same superfamily of cation channels as TPCs and is highly enriched in late endosomes and lysosomes, but not in other endomembrane organelles (*35, 36*). In contrast to TPC2, TRPML1 current densities were unchanged in CCZ1 KO cells compared with WT cells (Fig. S3C–E). These data suggest that CCZ1 loss selectively affects TPC2 rather than broadly altering endolysosomal cation channel activity. In addition to increasing maximal current amplitudes, CCZ1 deletion also enhanced TPC2’s apparent affinity for the agonist PI(3,5)P₂ by approximately 2.5-fold [EC50_WT_: 0.6779 μM, EC50_CCZ1KO_: 0.2743 μM] (Fig. 2I–K, Fig. S4D). Finally, TPC2 currents activated by >10 µM PI(3,5)P₂ in CCZ1 KO cells exhibited voltage-dependent adaptation at negative membrane potentials between -70 to -100mV (compare current traces in Fig. 2I and J, Fig. S5H-L) A similar phenomenon was observed in WT cells co-expressing TPC2 and RAB7a, suggesting that this effect reflects an intrinsic property of the fully activated channel rather than a specific consequence of CCZ1 deletion (shape of current trace in Fig. 2D, Fig. S5F-G).

To exclude the possibility that the pronounced effects observed following CCZ1 deletion resulted from off-target activity of the CRISPR-Cas9 system, we conducted a rescue experiment. CCZ1 was reintroduced into knockout cells via an expression vector, and the cells were subsequently analyzed (Fig. 3). In these rescued cells, the interaction between RAB7a and TPC2 was fully restored reaching levels comparable to those observed in WT cells (Fig. 3A, B, Fig S2E). Moreover, current densities in the rescued cells were similar to those of WT cells (Fig. 3C-E). To exclude apilimod-specific effects, we tested the RAB7a–TPC2 interaction in WT and CCZ1 KO cells without apilimod treatment and observed the same interaction pattern as with apilimod (Fig. S6). In agreement with previous findings (*26*) MON1a/b and RMC1 expression was decreased in CCZ1 KO cells (Fig. 3F). Importantly, upon re-expression of CCZ1, levels of both proteins were normalized (Fig. 3F).

**Figure 3:**
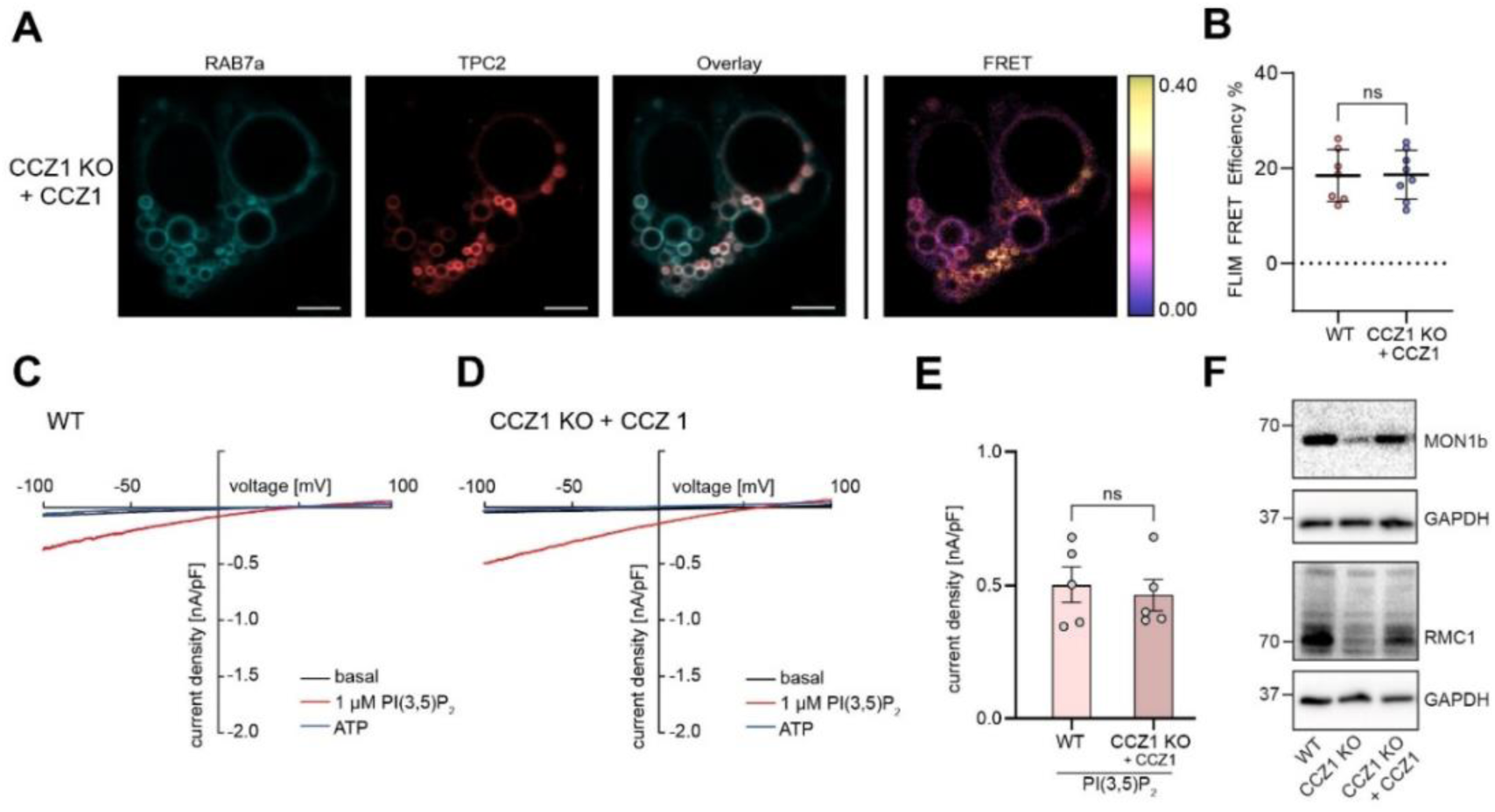
Re-expression of CCZ1 in HEK293 CCZ1 KO cells rescues RAB7A-TPC2 interaction and reduces TPC2 currents to WT levels. (**A**) Co-expression of mTq2-RAB7a (cyan), TPC2-mVenus (red) and CCZ1-mCherry in HEK293 CCZ1 KO cells, treated overnight with apilimod (1 µM), with corresponding FLIM-FRET efficiency maps shown on the right. FRET efficiencies (0–40%) are color-coded according to the color scale. Scalebar: 5 µm. (**B**) FRET efficiencies measured in representative regions of interest: representative FRET-positive vesicles in WT cells and representative FRET-positive vesicles in CCZ1 rescued CCZ1 KO cells (n=7 WT, from Fig. 2B, n=8 CCZ1 rescued CCZ1 KO cells from A). Error bars: ±SD. Statistical significance: Unpaired two tailed t-test, ns: not significant. (**C-D**) Whole-endolysosomal patch-clamp recordings of overexpressed TPC2 currents in WT (C) and in CCZ1 KO cells co-expressing CCZ1-mCherry plasmid (D), showing basal (black), ATP (blue), and PI(3,5)P₂ (red)-stimulated current densities. (**E**) Quantification of PI(3,5)P_2_-mediated TPC2 current densities at -100 mV from EL recordings in CCZ1 KO cells co-expressing CCZ1-mCherry plasmid (right bar), compared to WT cells (left bar). Data are presented as mean ± SEM. Statistical significance: unpaired two tailed t-test. ns, not significant. Each dot represents a single current density value measured from one EL, n=5 for each condition. (**F**) Representative Western blot of HEK293 WT, CCZ1 KO and CCZ1 KO transfected with CCZ1-expressing plasmid showing restored MON1B and RMC1 expression following CCZ1 transfection.

Taken together, these experiments confirm that the previously described effects were indeed caused by the absence of CCZ1 and are not attributable to non-specific alterations in the HEK293 genome.

### Constitutively active RAB7a^Q67L^ mutant does not rescue RAB7a function in the absence of CCZ1

CCZ1 is an essential component of the GEF complex that facilitates the exchange of GDP for GTP on RAB7a, thereby activating the protein. It is therefore conceivable that the impaired regulation of TPC2 activity in CCZ1 KO cells results from a failure to activate RAB7a. To explore this hypothesis, we co-expressed TPC2 with a constitutively active RAB7a mutant (RAB7a^Q67L^), which has been used previously to bypass GEF-function of CCZ1 in late endosomal trafficking and LDL-cholesterol uptake (*26*). As expected, RAB7a^Q67L^ showed strong interaction with TPC2 in WT cells (Fig. 4A, upper panel, Fig. 4B). However, there was no interaction in CCZ1-deficient cells (Fig. 4A, lower panel, Fig 4B). Consistent with this observation, RAB7a^Q67L^—like RAB7a—failed to stimulate TPC2 currents in CCZ1 KO cells (Fig.4C-E).

**Figure 4:**
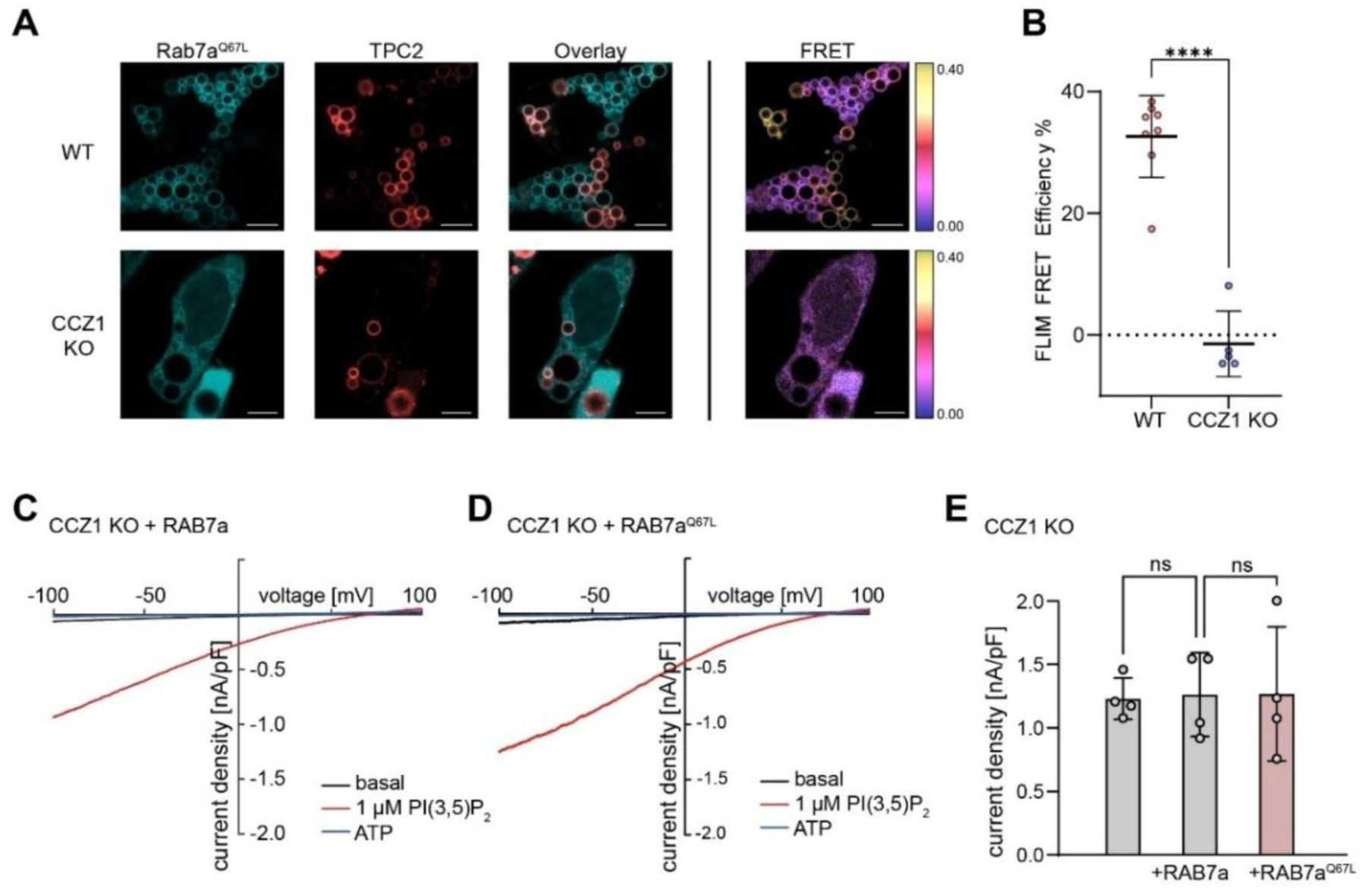
RAB7a^Q67L^ fails to interact with TPC2 and modulate its currents in CCZ1 KO cells. **(A)** Co-expression of mTq2-RAB7a^Q67L^ (cyan), TPC2-mVenus (red) in HEK293 WT and CCZ1 KO cells, treated overnight with apilimod (1 µM), with corresponding FLIM-FRET efficiency maps shown on the right. FRET efficiencies (0–40%) are color-coded according to the color scale. Scale bars: 5 µm. (**B**) FRET efficiencies measured in representative regions of interest (FRET-positive vesicles in WT cells and representative mTq2-RAB7a^Q67L^-positive areas in CCZ1 KO cells) corresponding to images in A (n=8 WT, n=5 CCZ1 KO). Error bars: ±SD. Statistical significance: Unpaired two tailed t-test, ****p<0.0001. (**C-D**) Whole-endolysosomal patch-clamp recordings of overexpressed TPC2 currents in CCZ1 KO cells with co-expression of RAB7a (C) or RAB7a^Q67L^ (D), showing basal (black), ATP (blue), and PI(3,5)P₂ (red)-stimulated current densities. (**E**) Quantification of PI(3,5)P_2_-mediated TPC2 current densities at -100 mV from EL recordings in CCZ1 KO cells co-expressing TPC2 with RAB7a^Q67L^ (pink bar) compared to cells with co-expressing RAB7a (middle grey bar) from Figure 2H. Data are presented as mean ± SEM. Statistical significance: one-way ANOVA with Bonferroni post-hoc tests. ns, not significant. Each dot represents a single current density value measured from one EL. ns: not significant, n=4 for each condition.

### The CCZ1-dependent regulation of TPC2 is not attributable to impaired endosomal maturation

Recent studies have shown that, in addition to its function as a RAB7a-GEF, the CCZ1-MON1a/b-RMC1 complex also recruits the GTPase-activating protein (GAP) TBC1D18 (also known as RABGAP1L) (*37*), which inactivates RAB5 and thus contributes to the RAB5-to-RAB7 conversion during the early-to-late endosome transition (*38, 39*). Enlarged endosomal compartments in CCZ1 KO cells displayed a late-endosomal signature (LAMP1⁺, TPC2⁺, EEA1⁻, Fig. 1F–H) and may arise from increased homotypic fusion due to impaired RAB5 inactivation. Supporting this, we observed increased fluorescence signal correlation of TPC2 with RAB5 in CCZ1 KO cells compared to WT (Fig. 5A, B). However, FRET efficiency remained at baseline levels in both cell types (Fig. 5C, D), indicating no direct interaction between TPC2 and RAB5.

**Figure 5:**
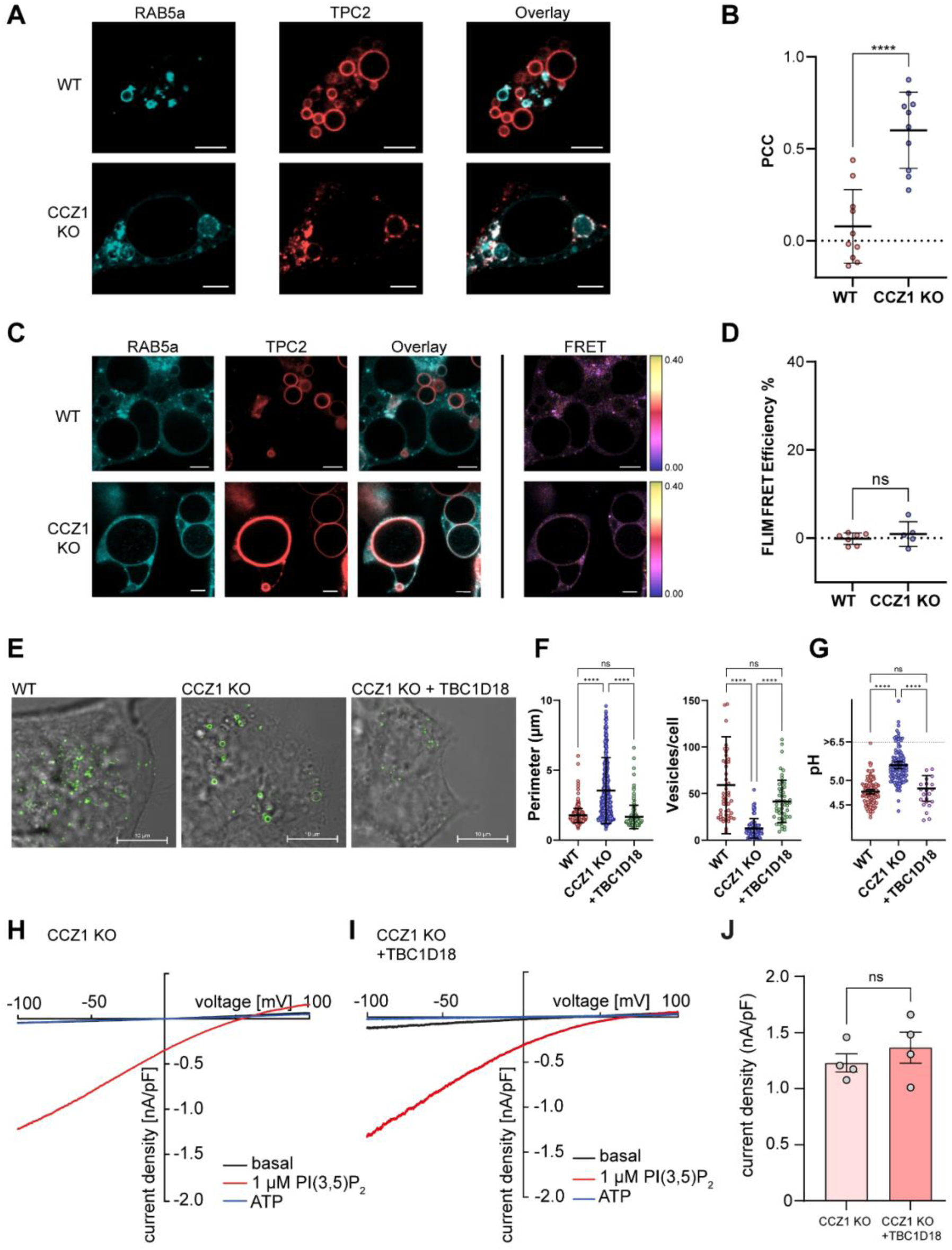
TBC1D18 decreases endolysosomal vesicle size in CCZ1 KO cells to WT level without affecting TPC2 activity. (**A**) Representative confocal images of WT and CCZ1 KO cells co-expressing mTq2-RAB5 (cyan) and TPC2-mVenus (red) treated with apilimod (1 µM). (**B**) Quantification of Pearson’s correlation coefficient between TPC2 and RAB5 signals from multiple cells. Each dot represents an individual cell (n=10 WT, n=10 CCZ1 KO); mean ± SD is indicated. Statistical significance: unpaired t-test, ****p<0.0001. (**C**) HEK293 WT (top row) and CCZ1 KO cells co-expressing mTq2-RAB5 (cyan) and TPC2-mVenus (red), treated with apilimod (1 µM), with corresponding FLIM-FRET efficiency maps shown on the right. FRET efficiencies (0–40%) are color-coded according to the color scale. Scalebar: 5 µm (**D**) FRET efficiencies measured in representative regions of interest (FRET-positive vesicles in WT cells and representative RAB5-positive areas in KO cells) corresponding to images in C (n=7 WT, n=5 CCZ1 KO). Error bars: ±SD. Statistical significance: Unpaired two tailed t-test, ns: not significant. (**E**) Confocal super-resolution (AiryScan) images of TPC2-mVenus overexpressing HEK293 WT (left image), CCZ1 KO cells (center image), and CCZ1 KO cells additionally overexpressing TBC1D18-HA (right image). (**F**) Left diagram: Quantification of the perimeter of TPC2-mVenus–positive endolysosomal membranes in WT and CCZ1 KO cells overexpressing TPC2-mVenus (WT: n=451, CCZ1 KO: n=496) and in CCZ1 KO cells co-expressing TPC2-mVenus and TBC1D18-HA (n=176), according to images in F. Right diagram: Quantification of TPC2-positive vesicles per cell in WT (n=49 cells), CCZ1 KO (n=73 cells) and CCZ1 KO cells co-expressing TBC1D18-HA (n=52 cells). Error bars for both diagrams: ±SD. Statistical significance of both diagrams: Kruskal Wallis test, ****p<0.0001, ns: not significant. (**G**) Ratiometric pH measurements of WT (n=124 vesicles, n=41 cells), CCZ1 KO (n=117 vesicles, n=64 cells), and CCZ1 KO cells overexpressing TBC1D18 (n=25 vesicles, n=11 cells). Each dot represents one endosome. Error bars: ±SEM. Statistical significance: Kruskal Wallis test, ****p<0.0001, ns: not significant. (**H-I**) Whole-endolysosomal patch-clamp recordings of overexpressed TPC2 currents in CCZ1 KO cells (H), in CCZ1 KO cells co-expressing of TBC1D18-HA (I), showing basal (black), ATP (blue), and PI(3,5)P_2_ (red)-stimulated current densities. (**J**) Quantification of PI(3,5)P_2_ mediated TPC2 current densities at -100 mV from CCZ1 KO cells with (right bar) or without co-expressing of TBC1D18-HA (left bar) from recordings Fig. 2H and 5I. Each dot represents a single current density value measured from one EL. Data are presented as mean ± SEM. Statistical significance: unpaired two tailed t-test. ns: not significant, n=4 for each condition.

Since overexpression of TBC1D18 has recently been shown to compensate for the lack of CCZ1 in RAB5 inactivation (*38*), we applied this approach to dissect CCZ1’s role as a RAB7a-GEF from its function in RAB5 regulation. Following transfection with HA-tagged TBC1D18, CCZ1 KO cells exhibited a strong band at the expected molecular weight of 92 kDa (Fig. S7A). Importantly, overexpression of TBC1D18 significantly reduced the size of TPC2-positive organelles (Fig. 5F, G), and their number increased back to WT levels. Because loss of RAB5 alone is insufficient to confer late endosomal identity or function, we assessed endosomal acidification as a functional readout of late endosome formation (Fig. 5G). In CCZ1 KO cells, endosomal acidification was arrested at approximately pH 5.5, consistent with compartments at or beyond the early-to-late endosomal transition. Notably, overexpression of TBC1D18 restored endosomal acidification to WT levels (pH ∼4.5–5), indicating functional recovery of late endosome/lysosome acidification and identifying RAB5 inactivation as a key rate-limiting step in endosomal maturation in CCZ1 KO cells. However, endolysosomal patch-clamp experiments revealed that this rescue had no effect on TPC2-mediated currents, which remained enlarged in CCZ1 KO cells, regardless of TBC1D18 overexpression (Fig. 5H–J). Furthermore, TBC1D18 did not restore the interaction between RAB7a^Q67L^ and TPC2 (Fig. S7B).

Taken together, our findings strongly suggest that the regulation of TPC2 by CCZ1 involves a mechanism that is independent of endosomal maturation and RAB5 activity.

### Gain of TPC2 function promotes enhanced migration in a CCZ1-deficient melanoma cell line

TPC2 has been implicated in the formation and progression of cancer cell migration, invasion, proliferation (*10, 11, 13, 24, 40*). The human melanoma cell line SK-MEL-5 endogenously expresses TPC2 and has been previously used as a model system to investigate the role of TPC2 and RAB7a in cancer cell migration (*24*). To extend our findings from HEK293 cells, we generated a CCZ1 KO in SK-MEL-5 cells (Fig. 6A, Fig. S1C). TPC2 was activated using the cell-permeable agonist A1P, which mimics PI(3,5)P₂ and was also previously used in patch-clamp and cell migration experiments (*24, 41*). Consistent with our previous results using PI(3,5)P_2_, patch-clamp recordings revealed that A1P-mediated TPC2 current densities were significantly increased in CCZ1 KO cells compared to SK-MEL5 WT cells. Normalized current densities were 0.92 ± 0.16 nA/pF (n = 4) in WT and 2.03 ± 0.07 nA/pF (n = 4) in CCZ1 KO cells (Fig. 6B–D). A similar increase using A1P was observed in CCZ1 KO HEK 293 cells relative to WT (Fig. S4A-C).

**Figure 6:**
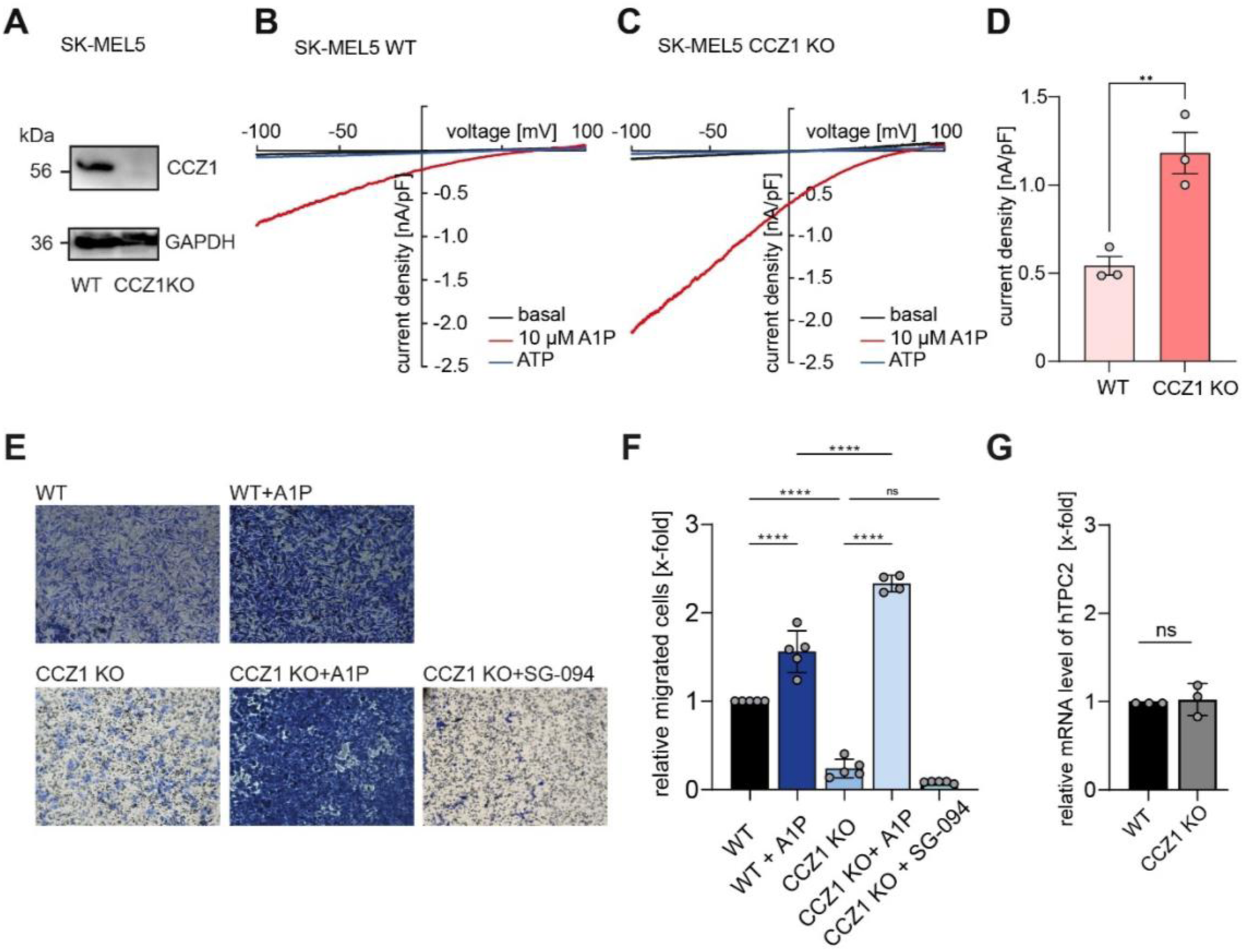
CCZ1 KO SK-MEL5 cells show enhanced TPC2 dependent migration. (**A**) Western blot validation of CRISPR-Cas 9 mediated CCZ1 deletion in SK-MEL5 cell line. (**B-C**) Whole-endolysosomal patch-clamp recordings of overexpressed TPC2 currents in WT (B) and CCZ1 KO (C) SK-MEL5 cells, showing basal (black), ATP (blue) and A1P (red)-stimulated current. (**D**) Quantification of A1P mediated TPC2 current densities at -100 mV from endolysosomal recordings from B and C. Each dot represents a single current density value measured from one EL. Data are presented as mean ± SEM. Statistical significance: unpaired two tailed t-test. **p<0.01 (p=0.0027), n=3 for each condition. (**E**) Representative images of 24h migration assay in SK-MEL-5 WT and CCZ1 KO cells, untreated or with treatment of A1P or SG-094, to specifically active or inhibit TPC2. Migrated cells were afterwards fixed and stained with crystal violet. (**F**) Total migrated cells from (E) were counted and normalized to WT. Each dot represents one independent experiment (one transwell). Data are presented as mean ± SEM. Statistical significance was determined using two-way ANOVA with Bonferroni post-hoc tests. ****p<0.0001, ns: not significant, n ≥4 for each condition. (**G**) Relative mRNAexpression of TPC2 in SK-MEL5 WT, CCZ1 KO cells, qPCR results were normalized by WT results, statistical significance was determined using unpaired two tailed t-test, ns: not significant, n=3 for each condition.

For cell migration assessment, we employed a Boyden chamber assay. Stimulation with A1P increased migratory capacity relative to unstimulated controls (Fig. 6E, F), with a notably greater effect observed in CCZ1 KO cells. Also, A1P stimulation resulted in significantly enhanced migration in CCZ1 KO cells compared to WT cells. This enhanced migration could be directly attributed to increased TPC2 activity, as it was completely abolished by treatment with the TPC2 inhibitor SG-094 (Fig. 6F) (*8, 42*). We also tested expression levels of MITF and GSK-3ß that were previously identified as downstream mediators coupling TPC2 activity with melanoma cell migration (*24*) (Fig. S9). As expected, GSK-3β levels were higher in KO cells compared to WT under basal condition while activation with A1P reduced GSK-3ß in KO cells to WT levels (Fig. S9A). This is consistent with a significant increase of migration in KO cells observed under these conditions. Similarly, reduced MITF levels of KO cells were normalized to WT levels upon TPC2 activation with A1P (Fig. S9B). Importantly, quantitative PCR analysis showed that TPC2 transcript levels were comparable between WT and CCZ1 KO SK-MEL cells, indicating that the absence of CCZ1 did not affect TPC2 expression itself (Fig. 6G).

## Discussion

Recent work has demonstrated that the small GTPase RAB7a enhances the activity of the endolysosomal two-pore channel TPC2 (*24*). While early proteomic studies had already identified RAB7a as part of the TPC2 interactome—suggesting a direct interaction between the two proteins (*23*) — our current study reveals that this functional relationship is more complex than previously assumed. Specifically, we show that the RAB7a-dependent regulation of TPC2 requires the presence of CCZ1 regardless of its GEF function.

CCZ1 forms a complex with MON1a/b and RMC1, which is known to function as the GEF for RAB7a, accelerating GDP-GTP exchange and thereby activating RAB7a. Using endolysosomal patch-clamp recordings in combination with confocal imaging and FRET analysis, we confirmed the interaction between TPC2 and RAB7a in WT HEK293 cells and showed that it is specific. However, this interaction was completely abolished in CRISPR/Cas9-engineered CCZ1 KO cells. Strikingly, the TPC2–RAB7a interaction was also lost in CCZ1 KO cells expressing a constitutively active RAB7a mutant (RAB7a^Q67L^), which has been used by others to bypass the GEF function of the CCZ1–MON1a/b–RMC1 complex (*26*). Although RAB7a^Q67L^ was able to bind TPC2 in WT cells (this study; (*23, 24*)), we observed no interaction in CCZ1-deficient cells, either by confocal imaging or FRET analysis. These findings suggest that CCZ1 is indispensable for the TPC2–RAB7a interaction, independent of its role as a GEF.

Another striking observation was that TPC2 currents in CCZ1 KO cells, while insensitive to RAB7a, were substantially larger than in WT cells. This effect was independent of the activating ligand, as it was observed with both the natural ligand PI(3,5)P₂ and the synthetic agonist A1P. Current densities in CCZ1-deficient cells were comparable to those seen in RAB7a-activated WT cells. These currents also displayed a modest increase in apparent PI(3,5)P₂ affinity, as well as voltage-dependent adaptation at membrane potentials between -70mV to -100mV — a feature also observed in RAB7a-activated currents in WT cells. This suggests that such adaptation may reflect a property of fully activated TPC2 currents or of large current amplitudes, rather than a CCZ1-specific effect. Moreover, This adaptation is consistent with a previous report of ligand-tuned TPC2 gating (*43*) and suggests that intracellular effectors like CCZ1 and RAB7a may similarly modulate this voltage dependent adaptive behavior, underscoring the intrinsic plasticity of TPC2. Notably, re-expression of CCZ1 in these KO cells restored the interaction and current densities, indicating that the observed phenotype was not due to off-target effects of Cas9.

Beyond its GEF activity, the CCZ1–MON1a/b–RMC1 complex also acts as a GTPase-activating protein (GAP) for RAB5, thereby promoting the transition from early (RAB5+) to late (RABa7+) endosomes not only via activating RAB7a, but also by governing RAB5 clearance. When this process is impaired, early endosomes tend to undergo homotypic fusion, resulting in abnormally large organelles (*44*). Consistent with this, we observed enlarged endolysosomal compartments in CCZ1 KO cells, along with increased co-localization of TPC2 and RAB5.

Given this dual role of CCZ1, it is conceivable that the observed loss of RAB7a regulation could be secondary to disrupted endosomal maturation. However, we largely excluded this possibility using rescue experiments with TBC1D18, a recently identified mediator of the RAB5-GAP activity of the CCZ1 complex (*38*). Overexpression of TBC1D18 normalized both organelle size and number. Notably, the elevated intraorganellar pH of CCZ1 knockout endosomes was fully restored to WT levels (pH ≈ 4.5–5.0) upon TBC1D18 overexpression. Nevertheless, even under these “RAB5-GAP rescued” conditions, RAB7a^Q67L^ failed to interact with TPC2, nor did it modulate TPC2-mediated currents. These findings reveal a previously unrecognized function of the CCZ1 complex, independent of its known GEF and GAP activities, that facilitates RAB7a-mediated TPC2 regulation.

Taken together, our electrophysiological data are consistent with a model in which CCZ1 exerts an inhibitory influence on TPC2 that is lifted upon CCZ1 loss. Thus, RAB7a may activate TPC2 by functionally uncoupling it from the CCZ1–MON1a/b–RMC1 complex. The exact molecular mechanism remains to be elucidated. Our preliminary FRET experiments do not support a direct interaction between TPC2 and the trimeric complex. This could be due to steric hindrance imposed by the FRET tags, or alternatively, the regulation may involve additional, as-yet unidentified proteins acting as intermediaries between TPC2 and the CCZ1 complex. A working model based on our findings is highlighted in Figure 7.

**Figure 7.**
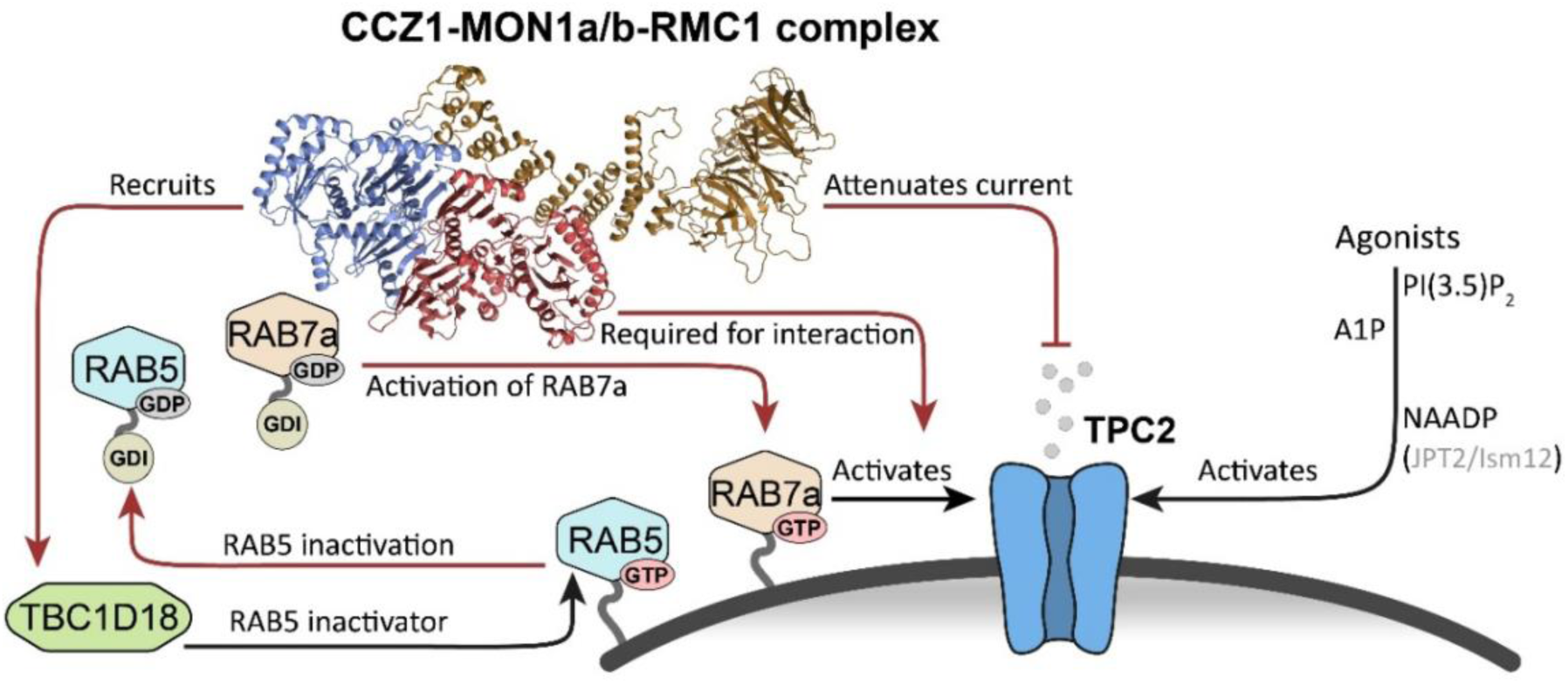
Working model summarizing CCZ1-MON1a/b-RMC1 functions in TPC2 regulation. The CCZ1/MON1/RMC1 complex is depicted using its cryo-EM structure (PDB: 9L0D,(*42*)). Red arrows denote CCZ1/MON1/RMC1-driven function: RAB7a activation, activation-independent promotion of RAB7a-TPC2 association, RAB5 inactivation, and attenuation of TPC2 currents. Inactive (GDI-bound) RAB5 and RAB7a are shown on the left; active, GTP-bound membrane-associated RAB5/RAB7a are shown on the endolysosomal membrane (gray surface). TPC2 is shown in blue, and the alternative RAB5 inactivator TBC1D18 in green.

Previous studies have established TPC2 as a critical regulator of cancer progression. In melanoma, TPC2 activity correlates with invasiveness: the channel is highly active in migratory cells, and pharmacological inhibition or genetic knockout slows migration (*9–11, 23*). Here, we identify CCZ1-mediated regulation of TPC2 as a novel mechanism controlling melanoma cell migration. In the SK-MEL-5 melanoma line, which endogenously expresses TPC2, CCZ1 exerted an inhibitory effect comparable to that observed in HEK293 cells. Consistently, CCZ1 knockout markedly enhanced migration. Deletion of CCZ1 also modulated downstream effectors of TPC2, including MITF and GSK-3β (Fig. S9), consistent with channel activation (*24*). Pharmacological inhibition of TPC2 suppressed migration irrespective of CCZ1 status. As a central regulator of RAB7a and RAB5, CCZ1 participates in multiple cellular pathways. It is therefore conceivable that the net effect on migration following CCZ1 deletion reflects a composite phenotype, incorporating TPC2 overactivity as well as additional defects arising from disrupted RAB7a-dependent processes.

Little is known about a role for CCZ1 in cancer. Although CCZ1 upregulation has been linked to progression of squamous cell carcinoma (*45*), it has not previously been implicated in melanoma. Analysis of public datasets reveals marked heterogeneity in CCZ1–MON1a/b–RMC1 expression across cutaneous melanoma cell lines (Fig. S10), consistent with context-dependent functions. Accordingly, the effects of CCZ1 modulation on cancer cell migration may reflect not only increased TPC2 activity but also perturbation of RAB7a- and RAB5-dependent trafficking pathways. Despite this complexity, our data identify CCZ1-complex status as a potential marker of melanoma contexts with altered RAB7a/TPC2 signaling and associated transcriptional and proliferative programs, although its clinical relevance remains to be established. Given the widespread expression of TPC2 and the diverse cellular processes in which this channel participates (*3, 35*), CCZ1-mediated regulation of TPC2 is likely to extend beyond oncogenic contexts and may have broader (patho)physiological relevance.

## Materials and Methods

### Plasmid generation

All ligations were done using T4 Ligase (ThermoFisher, Cat. EL0011). All constructs use pcDNA3.1+ as a backbone. N-terminally tagged mTq2-RAB7a was generated using an extension overlap PCR (Roche High Fidelity PCR Master, Cat. 12140314001) to combine mTurquoise2 and human RAB7a (CCDS3052.1). Insertion of the amplicon was done using restriction-ligation cloning in pcDNA3.1+ using NheI (ThermoFisher, Cat. FD0974) and HindIII (ThermoFisher, Cat. FD0504). mTq2-RAB7a^Q67L^ was generated using QuikChange Site-Directed Mutagenesis Kit (Agilent, Cat. 200519) with the following primers: 5ʹ-CTGGGGCGTGCTGTGCTTCGCCC-3ʹ, 5ʹ-GGGCGAAGCACAGCACGCCCCAG-3ʹ. mTq2-RAB5 was similarly generated by inserting mTq2 via NheI and BamHI (ThermoFisher, Cat. FD0054), and RAB5 (CCDS2633.1) via BamHI and ApaI (ThermoFisher, Cat. FD1414). C-terminally tagged TPC2-mVenus and TPC2-mTq2 (TPC2: CCDS8189.1) were generated similarly by human TPC2 insertion via NheI and EcoRI (ThermoFisher, Cat. FD0274), and Fluorophore insertion via EcoRI and XhoI (ThermoFisher, Cat. FD0694). LAMP1-mTq2 was generated by insertion of LAMP1 (CCDS41909.1) via HindIII and EcoRI into the C-terminal mTurquoise2 vector used before. TBC1D18 was synthesized de novo with a N-terminal HA tag separated from TBC1D18 by a Gly-Gly linker from TWIST Bioscience. CCZ1-mCherry was generated by insertion of CCZ1 (CCDS34597.1) via NheI and BamHI and mCherry via BamHI and ApaI. For patch-clamp measurements, the human TPC2-YFP plasmid (C-terminal tag) was used(*24*), and a mCherry-RAB7a plasmid that was generated by inserting mCherry via NheI and BamHI, and RAB7a via BamHI and ApaI. Calibration constructs for Two-Hybrid FRET assays were described previously(*33*). Designed sgRNA sequences targeting exon 10 (5ʹ-AAAGCACACAGCCGCACTCA-3ʹ) and exon 11 (5ʹ-CTTCGGCAAAAATCCAACGT-3ʹ) of human CCZ1 gene together with accessory sequences (U6 promoter sequence, scaffold sequence and RNA polymerase III start site sequence) were commercially purchased from IDT, inserted into Cas9 containing pcDNA3.1+ plasmid using Gibson assembly (ThermoFisher, Cat. A46624).

### Cell culture

Human embryonic kidney (HEK293) cells were maintained in DMEM (Invitrogen, #21885-025), supplemented with 10% fetal bovine serum (FBS; Thermo Fisher Scientific) and 100 U/mL penicillin–streptomycin (Pen/Strep; Sigma-Aldrich). SK-MEL-5 melanoma cells were cultured in high-glucose DMEM (Invitrogen, #31966-021) with 10% FBS and 100 U/mL Pen/Strep. All cell lines were cultured at 37 °C in a humidified incubator with 5% CO₂.

### CRISPR/Cas9 knockout lines

CRISPR/Cas9-mediated gene disruption of CCZ1 was performed in SK-MEL-5 melanoma and HEK293 cell lines using guide RNAs targeting exons 10 and 11 (see Fig. S1A-C) (*46*). sgRNAs were cloned into the pcDNA3.3_Cas9wt_Mali_sgRNA vector using Gibson assembly (ThermoFisher, Cat. A46624). Following transfection, single-cell clones were obtained by limiting dilution. Clonal populations were expanded and screened for successful knockout by genomic PCR and immunoblotting. For Western blot validation, an anti-CCZ1 antibody (B-7, Santa Cruz Biotechnology) was used.

### Western blotting

Cells were washed three times with ice-cold PBS and lysed in 400 µL lysis buffer (5M NaCl, 2.5M CaCl_2_ and 0.5% Triton X-100) supplemented with 1x protease inhibitor (Roche, Cat. 04693159001) per well. Following lysis, cells were scraped and transferred into Eppendorf tubes, then incubated on ice for 30 minutes. Lysates were centrifuged at 13,200 × g for 10 minutes at 4 °C, and the resulting supernatants were transferred to fresh tubes. Protein concentration was determined using the Qubit Protein Assay Kit (Thermo Fisher Scientific). Equal amounts of protein were mixed with 6× Laemmli sample buffer containing dithiothreitol (DTT) and heated at 72 °C for 10 minutes. Proteins were resolved by SDS–PAGE using self-cast gradient gels composed of 6% and 12% polyacrylamide. 15 µg of total protein per sample were loaded into wells, and molecular weights were estimated using the PageRuler Prestained Protein Ladder (Thermo Fisher Scientific). Electrophoresis was run at 100 V for 20 minutes, then at 150 V for 40 minutes. Proteins were transferred to methanol-activated PVDF membranes using a Mini Trans-Blot Cell (Bio-Rad) at 350 mA for 1.5 hours. Membranes were blocked for 1 hour at room temperature and incubated overnight at 4 °C with primary antibodies. Following three 10-minute wash steps, membranes were incubated with HRP-conjugated secondary antibodies for 2 hours at room temperature, followed by three additional washes. Chemiluminescent detection was performed using Western Blotting Luminol Reagent (Santa Cruz Biotechnology), applied according to the manufacturer’s protocol. Signal was visualized with a ChemiDoc MP Imaging System (Bio-Rad). The following antibodies were used with indicated dilution: CCZ1 (Santa Cruz, 1:1000, Cat. #Sc-514289), GAPDH (Cell-Signaling, 1:1000, Cat. #14c10), Anti-Mouse (Jackson, 1:1500, Cat. #715-035-150), Anti-HA (Cell Signaling, 1:1000, Cat. #6E2). Anti-MON1a (Proteintech, 1:1000, Cat. #23772-1-AP), Anti-MON1b (Proteintech, 1:1000, Cat. #17638-1-AP), Anti-MITF (cell Signaling, 1:1000, Cat. #97800), Anti-GSK-3ß (cell Signaling, 1:1000, Cat. #9832), Anti-H3 (cell Signaling, 1:1000, Cat. #9715),.

### Immunofluorescence staining

HEK293 WT and CCZ1 KO cells were seeded on 8 well removable chamber slides (Ibidi, Cat. #80841 with 1×10^5 cells per Well. On the following day, cells were washed 3x with PBS and incubated in freshly made 4% PFA (Thermofisher Scientific, Cat. #28906) for 10 min. at RT. Cells were then incubated for 20 min. at RT in 0.1 % Saponin solution (Sigma-Aldrich, Cat. #SAE0073), and afterwards blocked with 3 % BSA solution (Sigma-Aldrich, Cat. #A3059) for 1 h at RT. Primary Antibodies (RAB7a, Cell Signaling, #9367, 1:100; RAB5a, Cell signaling, #46449, 1:100; LAMP1, Cell Signaling, #9091S, 1:100; EEA1, Cell Signaling, #48453S, 1:100) were incubated overnight at 4° C. Cells were then washed 3x with PBS and incubated for 1h at RT with secondary Antibodies (Alexa Fluor 555, Invitrogen, #A1572, 1:500; Alexa Fluor 488, Cell Signaling, #89853, 1:500). Cells were then treated with Hoechst 33342 (Thermofisher Scientific, Cat. #H3570) according to the manufacturer. The wells were then removed from the slide and the mounting media Fluoromount-G (Invitrogen, Cat. #00-4958-02) added to the coverslip. Coverslips were sealed with transparent nailpolish and kept dark at 4° C until imaging.

### Whole-endolysosomal Patch-Clamp experiments

Endolysosomal patch-clamp experiments were conducted using a manual approach adapted from established protocols (*47*). WT and CCZ1 KO HEK293 and SK-MEL5 cells were plated on poly-L-lysine-coated glass coverslips (Sigma-Aldrich) in 24-well plates, maintaining an initial confluency of 60–70%. Transient transfection was carried out using TurboFect Transfection Reagent (Thermo Fisher Scientific) with plasmids encoding the target proteins. For co-expression of human TPC2 and RAB7a, a plasmid ratio of 2:1 was employed. To promote the formation of enlarged endolysosomal compartments, HEK293 cells were incubated with 1 μM apilimod (Axon Medchem) for overnight post-transfection. Residual apilimod was removed by washing prior to electrophysiological analysis. Patch-clamp measurements were performed at room temperature using an EPC-10 amplifier and PatchMaster software (HEKA Elektronik). Recording pipettes were refined to a tip resistance of 4–6 MΩ via fire polishing. Junction potentials were compensated electronically. Current signals were digitized at 49 kHz and filtered using a 2.8 kHz low-pass Bessel filter. Recordings were performed using voltage ramps (500 ms, –100 mV to +100 mV) applied at 5-second intervals. Current responses were quantified at –100 mV.The extracellular (bath) solution comprised 140 mM K-methanesulfonate (K-MSA), 5 mM KOH, 4 mM NaCl, 0.39 mM CaCl₂, 1 mM EGTA, and 10 mM HEPES, adjusted to pH 7.2 using KOH and to 300 mOsm with D-(+)-glucose. The pipette (luminal) solution contained 140 mM Na-MSA, 5 mM K-MSA, 2 mM Ca-MSA, 1 mM CaCl₂, 10 mM HEPES, and 10 mM MES, titrated to pH 4.6 with methanesulfonic acid and adjusted to 310 mOsm. Data were analyzed using OriginPro (OriginLab) and GraphPad Prism (GraphPad Software).

### FLIM FRET

HEK293 and HEK293 CCZ1 KO cells were seeded on ibidi 35 mm dishes with 1.5H glass bottom (Cat: 81158) for inverted microscopy, or ibidi channel slides (Cat: 80606) for upright microscopy depending on the microscope used. The FRET pair consisted of mTurquoise2 (*48*) and mVenus (*49*) and was transfected one day before the experiment using Lipofectamine 2000 (Thermo Fisher, Cat: 11668027) with 0.7 μg/ml donor (mTq2 linked to the respective Rab-protein) and 1.3 μg/ml acceptor (TPC2-mVenus). Optionally, cells were also treated with apilimod (1 μM) overnight. Single confocal sections were recorded at room temperature (RT). Images were acquired using time correlated single photon counting technique (TCSPC) (*50*). If not mentioned otherwise, experiments were conducted using a Leica STELLARIS 8 confocal microscope from Leica Microsystems (Mannheim, Germany), which was kindly made available by Leica Microsystems, based on an inverted DMi8 stand. The 448 nm line of a White Light Laser (WLL) pulsed at 40 MHz and a 63X Oil objective were used. Detection was with a HyD X Detector (hybrid photomultiplier detector) at 455-510 nm. Acceptor fluorescence was measured consecutively using a 514 nm laser and detection at 525-561 nm. Pinhole size was set to 95.6 µm (1 AU for 580 nm). FLIM images for Figure S6 were acquired using a commercial Leica SP8 WLL upright confocal microscope based on an upright DM8 stand. A 40 MHz pulsed 405 nm laser was used to excite the donor (mTurquoise2), detection was with a HyD SMD Detector at 450-490 nm. Acceptor fluorescence was measured consecutively using a 514 nm laser and detection at 520-580 nm. Pinhole size was set to 100.3 µm (1 AU for 580 nm). All images were acquired with a pixel size ranging from 65-262 nm with a minimum pixel dwell time of 1.4 µs. FLIM evaluation was done using LAS X v4.8.2. Donor lifetime was determined experimentally for Donor (mTurquoise2) alone and Donor fused to RAB7a. Donor lifetime values for both controls remained very close (3.9-4 ns) to previously reported values (*51*) on each instrument and were used as a baseline for each session. Absolute FRET values were determined via multi-exponential fits for fluorescence lifetime decay curves measured and calculated against the donor-only lifetime for each pixel. FRET values (0–40%) were visualized as heatmaps using a customized version of the Matplotlib ’magma’ colormap (low: R=71, G=53, B=153; mid: R=211, G=58, B=82; high: R=225, G=225, B=123).

### Two-Hybrid FRET

HEK293 and HEK293 CCZ1 KO cells were seeded on ibidi 35 mm dishes with 1.5H glass bottom (Cat: 81158). The same constructs were used for FLIM and Two-Hybrid FRET measurements. Single confocal sections were acquired using a commercial Leica SP8, based on a DMi8 stand at RT with a 40X Oil objective. Three fluorescence channels were set up: mTq2DIRECT (458 nm laser excitation, 462 nm - 493 nm detection via HyD Detector), mTq2FRET (458 nm laser excitation, 524 nm - 548 nm detection via HyD SMD Detector, captured simultaneously with mTq2DIRECT), mVenusDIRECT (514 nm laser excitation, 524 nm - 548 nm detection via HyD SMD Detector). Pinhole was set to 1.2 AU (65 µm, calculated for 485 nm). All images were acquired with a pixel size ranging from 79-148 nm and a minimum pixel dwell time of 0.8 µs. After acquisition, regions of interest were drawn around vesicles in Fiji. The mean fluorescence intensity in each channel was determined and exported as .csv files. Calibration constants were obtained from HEK293 cells expressing single mTq2, mVenus, and three tandem constructs combining mTq2 and mVenus with different linker lengths, individually (0.5 μg/ml transfected plasmid for each). Cells transfected with the FRET pair of interest were additionally treated with 1 μM apilimod overnight. Calculation and binding curve generation was done using a custom MatLab function (MathWorks) (*33*). Two-Hybrid FRET heatmaps were generated using a custom Fiji macro calculating absolute FRET efficiencies for each pixel and plotting them in the same color-scale used for FLIM maps (see section FLIM FRET).

### AiryScan confocal imaging

HEK293 and HEK293 CCZ1 KO cells were seeded on glass bottom ibidi channel slides (Cat: 80607). One day before the experiment, cells were transfected with 1.3 μg/ml TPC2-mVenus plasmid and optionally 2 μg/ml TBC1D18 plasmid. Images were acquired on a commercial ZEISS LSM980 AiryScan at RT. The AiryScan mode was set up with a 514 nm laser and an additional T-PMT channel. Pinhole was set to 5.19 AU for 417 nm. Pixel dwell time was 1.27 µs. TPC2-positive puncta were then classified in two categories: Halo-shaped fluorescent circles were classified as in-focus vesicles, while dot-like structures were considered slightly out-of-focus vesicles. Vesicle diameter evaluation was determined by redrawing all halo-shaped objects in Fiji measuring the perimeter. For the vesicle number/cell evaluation, all objects (halo-shaped and dot-like structures) were counted for each cell. Statistical analysis was done using Mann-Whitney test if 2 groups were compared, and Kruskal-Wallis test if 3 groups were compared.

### Co-localization experiments

HEK293 and HEK293 CCZ1 KO cells were seeded on ibidi channel slides (Cat: 80606). Proteins of interest were transfected using Lipofectamine 2000 (Thermo Fisher, Cat: 11668027) and optionally treated with 1 μM apilimod one day before imaging. Image acquisition was done on a commercial ZEISS LSM980 AiryScan upright confocal microscope at RT using single confocal sections. mTurquoise2 channel was acquired using a 445 nm laser and detection at 454-517 nm, the mVenus channel with a 514 nm laser and detection at 534-659 nm, a Pinhole of 63 μm (1 AU for 445 nm) and a pixel dwell time of 4.84 μs. Pixel size was 78 nm. All images were acquired in 16 Bit. A fixed (−500) background subtraction of all images treated with apilimod was conducted in Fiji. Threshold level for PCC calculation was set and identical for all images within an experimental group (e.g. WT and KO with TPC2+LAMP1) to include both organellar Donor and Acceptor fluorescence but not cytosolic regions. Statistical analysis was done using unpaired t-tests.

### Electron microscopy

HEK293 and HEK293 CCZ1 KO cells were fixed in cell culture dishes for 30 minutes in 150 mM HEPES at pH 7.4, containing 1.5 % formaldehyde and 1.5 % glutaraldehyde. The sample was then immobilized in 2 % agarose and incubated for 2 hr in 1 % OsO4 aqueous solution with 1.5 % hexacyanoferrat II. Samples were then washed with ddH2O and stored in 1 % aqueous uranyl acetate at 4°C overnight. Cells were embedded in Epon after washing with ddH2O and dehydration with acetone. Samples were sectioned in 60 nm slices and mounted on formvar-coated copper grids and poststained with uranyl acetate and lead citrate. Imaging was done on a Morgagni TEM using a Veleta CCD camera. Two independent cultures were imaged for both WT and KO cells.

### Ratiometric pH measurements

10mg/ml stocks of OregonGreen-Dextran 10k MW and AlexaFluor568-Dextran 10k MW were prepared in PBS. HEK293 and HEK293 CCZ1 KO cells were seeded on ibidi 35 mm dishes with 1.5H glass bottom (Cat: 81158). One day before the experiment, additional HEK293 CCZ1 KO cells were transfected with TBC1D18-HA+mTq2 (1 µg/mL). On the day of the experiment, 30 µL of both Dextran conjugates were added to 1 mL of pre-warmed medium, briefly inverted and used immediately to replace the growth medium on the cells. The Dextran-conjugates were loaded for 15 min., replaced with pre-warmed medium and chased for 20 min. Afterwards, cells were imaged immediately for a maximum of 5 min. to avoid further chasing during imaging. Each condition was loaded, chased and imaged individually. To detect endosomes automatically, a detection workflow was created in Fiji: The AlexaFluor568 image was used and a weak Gaussian Blur (1.5 px) applied. A global threshold for all images was set and the binary mask used to detect particles with a minimum of 0.1 µm^2^ area. Particles were then quantified in terms of integrated intensity for both the OregonGreen and AlexaFluor568 channel, the ratio of OG/AF568 integrated intensity calculated and used for pH assessment. pH calibration was conducted using WT cells with the same preparation and additional incubation with 10 µM Nigericin and 10 µM Monensin for 10 min. before image acquisition. Ratios for pH 7.5, 6.5, 5.5 and 4.5 were determined and plotted with a cubic spline curve to derive pH values for samples. pH 7.5 ratio was not included as it was similar to pH 6.5.

### RT-qPCR

Total RNA was extracted from cellular samples using the RNeasy Mini Kit (Qiagen) following the manufacturer’s protocol. After centrifugation, cells were washed twice with ice-cold PBS and resuspended in ice-cold RLT buffer supplemented with 40 µM DTT (R0861, Thermofisher). The concentration of the isolated mRNA was quantified using a Nanodrop Spectrophotometer. Subsequently, 2000 ng of RNA was reverse transcribed into cDNA using the High-Capacity cDNA Reverse Transcription Kit (Applied Biosystems, Waltham, MA, USA) in accordance with the supplier’s guidelines. Quantitative real-time PCR (RT-qPCR) was performed using PowerUp™ SYBR® Green Master Mix (Applied Biosystems) in a reaction mixture containing 2 µl cDNA (equivalent to 50 ng), 6.25 µl PowerUp™ SYBR® Green Master Mix, 3.25 µl RNase-free water, and 0.5 µl (200 nM) of both forward and reverse primers per well. The amplification reactions were carried out on the QuantStudio™ 3 Real-Time PCR System (Applied Biosystems), and relative gene expression levels were analyzed using the ΔΔCT method (*52*). ACTIN was used as a housekeeping gene. Primers were purchased from Metabion (Planegg, Germany) and validated for their specificity and efficiency prior to use. The sequences of primers used for RT-qPCR were: hTPCN2 (fw): 5ʹ-TGCATTGATCAGGCTGTGGT-3, hTPCN2 (rev): 5ʹ-GAAGCTCAAAGTCCGTTGGC-3ʹ, hACTIN (fw): 5ʹ-CCAACCGCGAGAAGATGA-3ʹ, hACTIN (rev): 5ʹ-CCAGAGGCGTACAGGGATAG-3ʹ.

### Migration assay

A total of 0.5 × 10⁶ WT or CCZ1 KO SK-MEL-5 cells were seeded into the upper chamber of Transwell inserts (8 μm pore size; Sigma-Aldrich) in serum-free medium. The lower wells of a 24-well plate were filled with culture medium supplemented with 10% FBS as a chemoattractant. For treatment conditions, both the upper and lower chambers were supplemented with 10µM SG094 and 20 µM A1P. Cells were incubated for 24 h at 37 °C. After incubation, non-migrated cells on the upper membrane surface were removed and migrated cells on lower side were fixed (4% PFA, 10min RT; Sigma Aldrich; 252549-25ML) and stained using crystal violet (Sigma Aldrich, V5265-250ML). Membranes were imaged using an EVOS M7000 microscope (Thermo Fisher Scientific), and migrated cells were quantified using ImageJ software (NIH, Bethesda, MD, USA).

## Statistical analysis

For patch-clamp experiments, analysis of sweeps was completed in Origin. For each condition at least three independent samples/recordings were included. Statistical analyses of patch-clamp recordings, FRET quantification, co-localization, vesicle perimeter, vesicles number and vesicle pH were performed in GraphPad Prism. All error bars were shown as ± SEM or ± SD, as specified in the figure legends. For comparisons between two independent groups, a two-tailed unpaired Student’s t-test was used for normally distributed data; otherwise, a two-tailed Mann–Whitney U test was used. For comparisons of >2 independent groups, one-way ANOVA was used for normally distributed data followed by post-hoc a Bonferroni test, whereas non-normally distributed datasets were analyzed by Kruskal–Wallis test followed by Dunn’s multiple comparisons test with multiplicity-adjusted P values. For experiments with two factors, two-way ANOVA was performed with a Bonferroni test. Exact (adjusted where applicable) P values are reported for key comparisons.

## Acknowledgments

We thank Dr. Marco Keller for kindly providing the compounds TPC2-A1-P and SG-094 used in our experiments. We also thank Mariano G. Pisfil and Steffen Dietzel at BioMedical Center, Ludwig-Maximilians-Universität, Munich, Germany, for their support in imaging and analysis. This work was supported by funding of the German Research Foundation (SFB/TRR152 P12 to M.B and S.M., P04 to C.G. and P28 and INST 192/543-1 FUGG to C.W.-S. C.-C.C. was supported by the National Science and Technology Council (NSTC 114-2320-B-002 -022 -MY3 and NSTC-DAAD 114-2927-I-002-509), National Taiwan University (NTU-111L7826) and National Health Research Institutes (NHRI-EX111-11119SC).

## Author Contributions

Z.Y, C.F, L.O. A.C.L, S.F. and C.-C.C. designed experiments and collected and analyzed data. Z.Y. A.C.L and C.-C.C. performed endolysosomal patch-clamp experiments. C.F. performed confocal imaging and FRET experiments. C.F. and M.S. performed electron microscopy imaging. K.B. and G.C. provided SK-MEL-5 cell line. Z.Y. and L.O. performed and analyzed migration assays. M.B., S.M., C.W.-S. and C.G. provided funding and commented on the manuscript. M.B., C.W-S. and C.-C.C. coordinated research, designed the study, and wrote the manuscript. All of the authors discussed the results and commented on the manuscript.

