## Supplementary information for "CCZ1 is a modulator of TPC2 activity and melanoma cell migration"

**This PDF file includes:**

Figures S1 to S10

**Fig. S1**

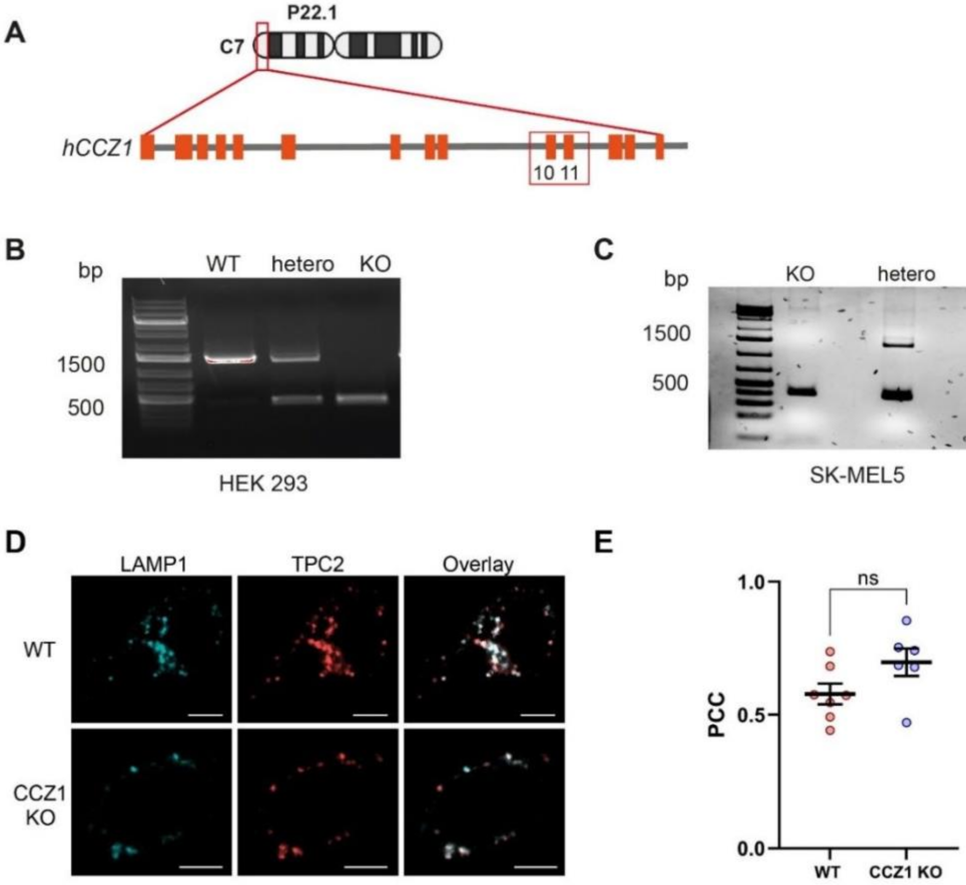


**Supplementary Figure S1**: **Validation of CCZ1 KO in HEK293 and SK-MEL-5 cells and apilimod control.** (**A**) Generation of the CCZ1 KO cell line using CRISPR-Cas9 targeting exon 10 and exon 11. Agarose gels show CCZ1 KO clone selection in HEK293 (**B**) and SK-MEL5 (**C**) PCR results of generated cell lines. Clones with a deleted 500 bp amplicon were used for Western blot. (**D, E**) Co-localization of lateendosomal marker protein with TPC2 without apilimod treatment. (**D**) Representative confocal images of WT and CCZ1 KO cells co-expressing LAMP1-mVenus (cyan) and TPC2-mTq2 (red). (**E**) Quantification of Pearson's correlation coefficient between TPC2 and LAMP1 signals in multiple cells as shown in A. Each dot represents an individual cell (n=10 WT, n=10 CCZ1 KO); mean ± SD is indicated. Statistical significance was assessed using an unpaired t-test, ns: not significant.

**Fig. S2**

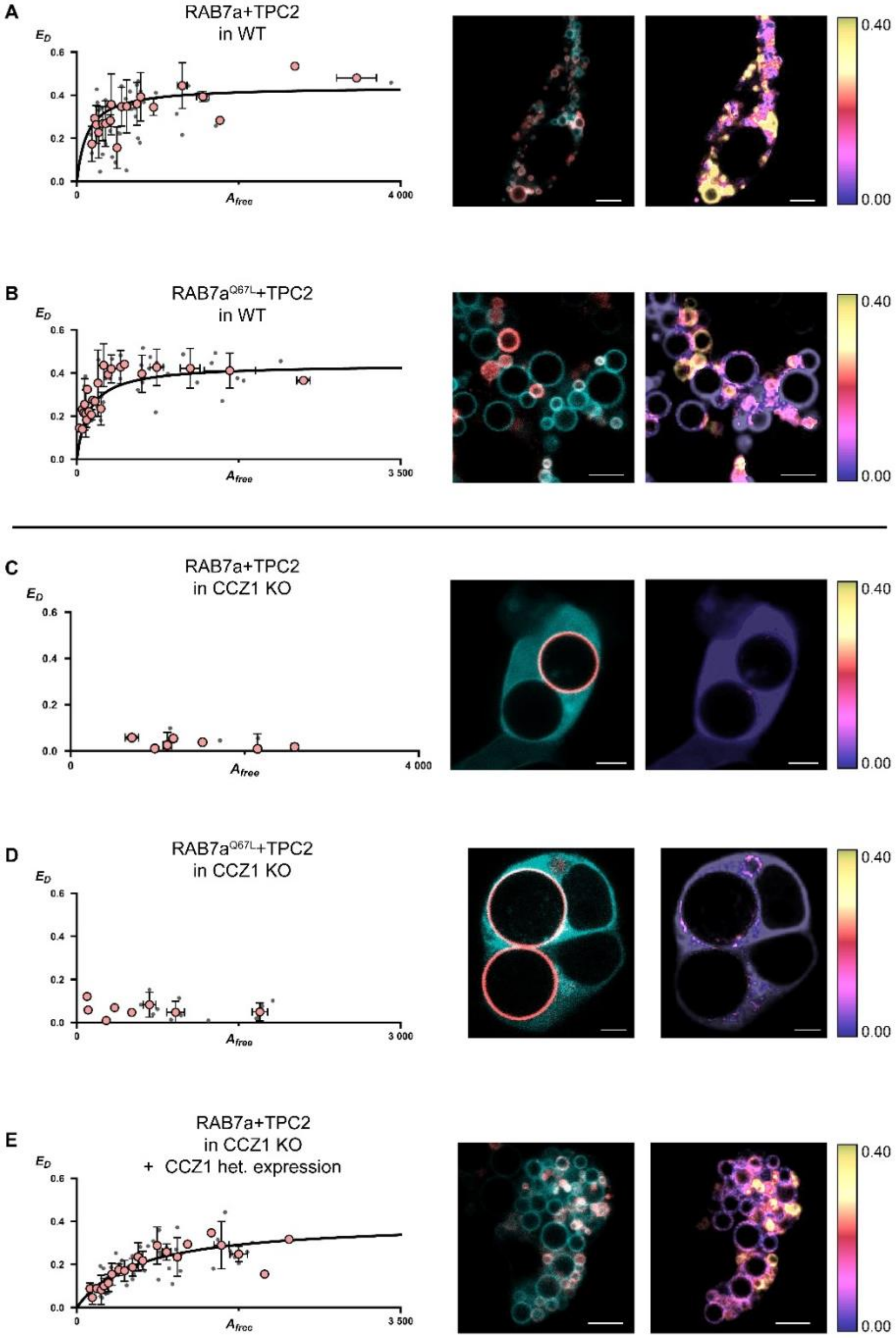


**Supplementary Figure S2: Two-Hybrid FRET assays were performed as a complementary FRET based approach to validate the interaction results obtained by FLIM-FRET.** Left diagrams: Two-Hybrid FRET binding curves of indicated protein-protein interactions. For each experiment, cells were co-transfected with the corresponding proteins (mTq2-RAB7a or mTq2-RAB7a[QL] and TPC2-mVenus) and treated with apilimod (1 µM). Y-axis (ED) indicates absolute FRET efficiency values calculated from FRET-related donor-quenching (mTq2-RAB7a). The X-axis indicates the estimated amount of free acceptor molecules (TPC2-mVenus). Each grey dot represents one region of interest, if possible, drawn at donor- and acceptor co-localizing vesicular structures (A: n=71 B: n=55, C: n=27, D: n=34 E: n=86, each binding curve: n=>15 different cells). Red dots represent binned data values with ±SD On the right, representative confocal images of each protein–protein interaction pair are shown, along with corresponding FRET efficiency (ED) maps calculated using Two-Hybrid FRET on a pixel-by-pixel basis. FRET efficiencies are color-coded according to the scale bar on the right. Scale bars: 5 μm.

**Fig. S3**

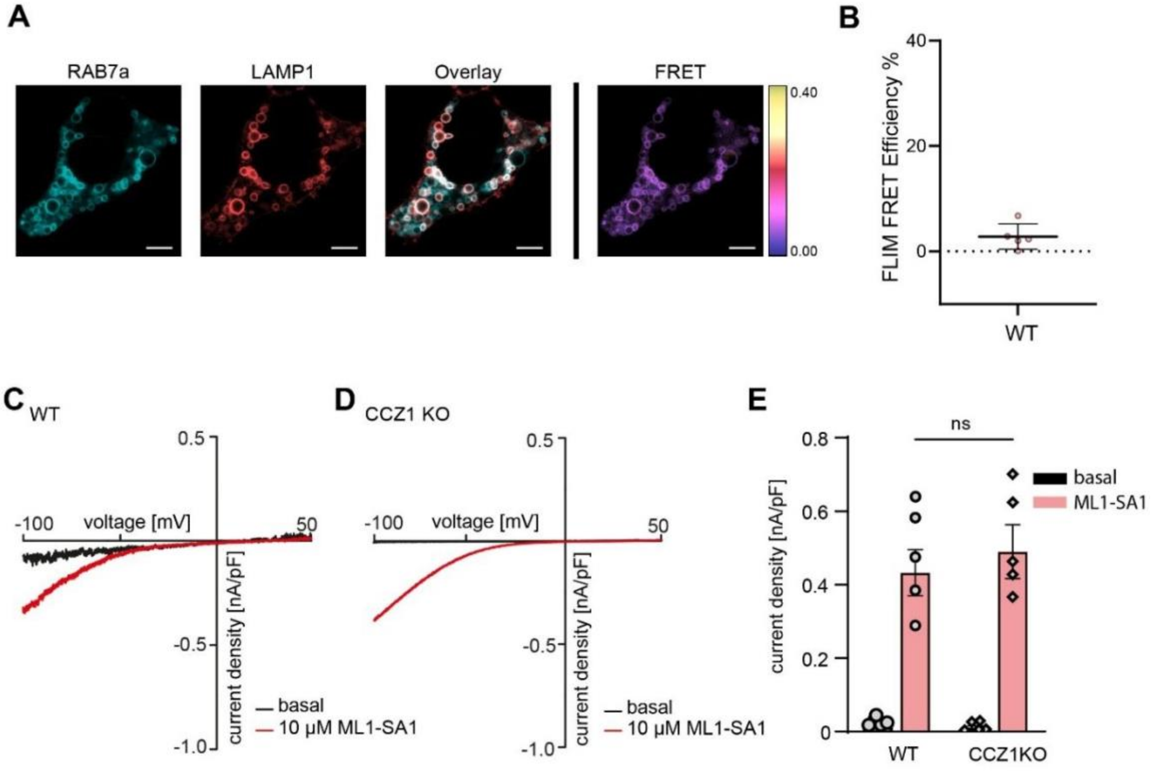


**Supplementary Figure S3: RAB7a-TPC2 FRET signals are specific and TRPML1 current densities are not affected by loss of CCZ1.** (**A**) HEK293 WT cells co-expressing mTq2-RAB7a (cyan) and Lamp1mVenus (red), treated with apilimod (1 µM), with corresponding FLIM-FRET efficiency map shown on the right. FRET efficiencies (0–40%) are color-coded according to the color scale. Scale bars: 5 µm. (**B**) FRET efficiencies at endosomal membranes with high levels of co-localization between Rab7 and Lamp1 (n=5). Error bars: ±SD. Statistical significance: Unpaired two tailed t-test, ****p<0.0001. (**C-D**) Wholeendolysosomal patch-clamp recordings endogenous TRPML1 currents in HEK293 WT and CCZ1 KO cells, showing basal (black) and ML1-SA1 (red)-stimulated current densities. Cells were pretreated with apilimod (1 µM) overnight. (**E**) Quantification of ML1-SA1-mediated TRPML1 current densities at -100 mV from endolysosomal recordings in (C, D). Data are presented as mean ± SEM. Each dot on the bar graph represents a single current density value measured from one endolysosome (EL). Statistical significance: unpaired two tailed t-test. ns, not significant.

**Fig. S4**

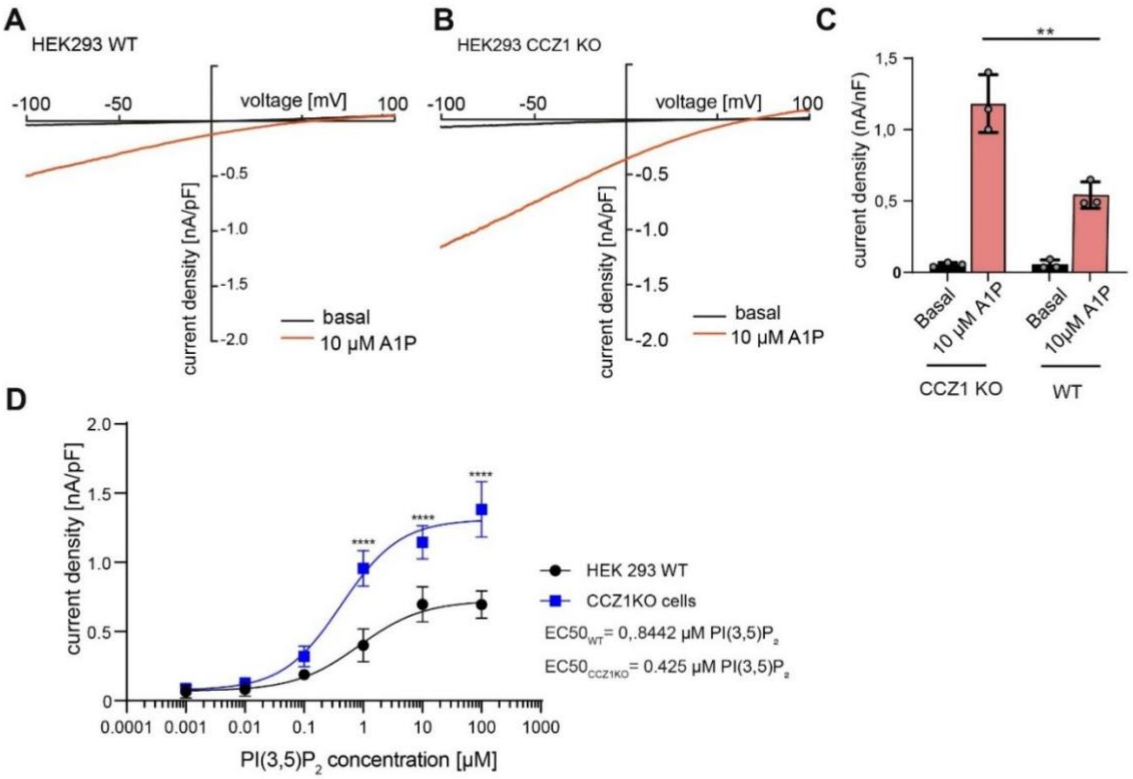


**Supplementary Figure S4: CCZ1 KO HEK293 cells showed a stronger current response to agonists compared with HEK293 WT cells.** (**A-B**) Whole-endolysosomal patch-clamp recordings of overexpressed TPC2 currents in WT (A) and CCZ1 KO (B) HEK293 cells, showing basal (black) and A1P (red)-stimulated currents. (**C**) Quantification of A1P mediated TPC2 current densities at -100 mV from endolysosomal recordings from A and B. Each dot represents a single current density value measured from one EL. Data are presented as mean ± SEM. Statistical significance was determined using unpaired two tailed t-test. **$p<0.01$. (**D**) Dose-response curve of overexpressed TPC2 comparing the current densities at -50mV induced by increasing concentrations of $PI(3,5)P_2$ in WT (black) and CCZ1 KO (blue) cells. Data are presented as mean ± SEM, statistical significance was determined using unpaired t-test (two-tailed), ****$p<0.0001$, n=4 for each condition.

**Fig. S5**

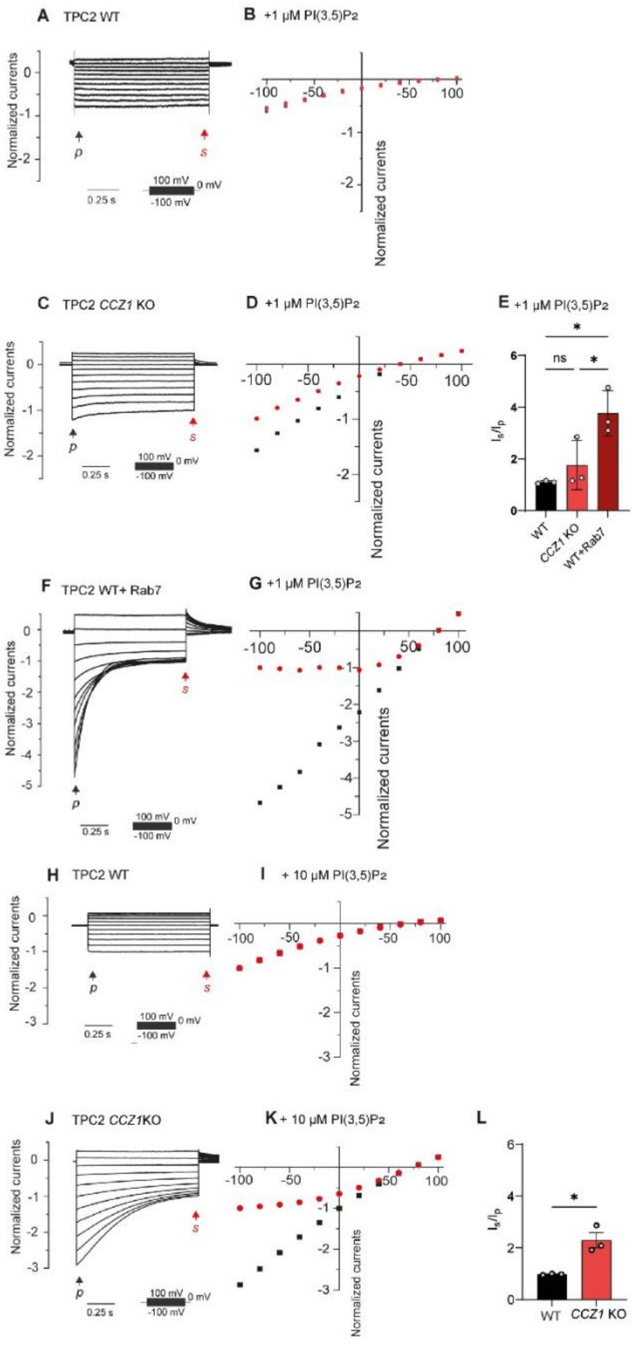


**Supplementary Figure S5: Voltage dependent adaptation of TPC2 at high polarized membrane potential.** (A, C, F) Whole-endolysosomal patch-clamp recordings showing normalized overexpressed TPC2 currents in HEK293 WT cells (A), in CCZ1 KO cells (C) and in HEK 293 WT with co-expression of RAB7a (F) in response to 1 µM PI(3,5)$P_2$, recorded using a step pulse protocol shown below the current traces. (B, D, G) Voltage-current (I-V) relationships for peak (p, red) and steady-state (s, black) currents in WT cells (B), in CCZ1 KO cells (D) and in WT with co-expression of RAB7a (G) with treatment of 1 µM PI(3,5)$P_2$. (E) Quantification of the $I_p/I_s$ ratio in WT (black) and CCZ1 KO (hell red) and WT with RAB7a co-expression (dark red) from A, C and F. Data are presented as mean ± SEM, *P < 0.05 (unpaired t-test). (H, J) Normalized overexpressed TPC2 currents in HEK 293 (H) and CCZ1 KO cells (J) treated with 10 μM PI(3,5)$P_2$, recorded using a step pulse protocol. (I, K) Voltage current (I-V) relationships for peak (p, red) and steady-state (s, black) currents in HEK 293 WT (I) and in CCZ1 KO (K) cells in response to 10 μM PI(3,5)P2. (L) Quantification of the $I_p/I_s$ ratio in WT (green) and CCZ1 KO (hell red) cells from H and J in response to 10 μM PI(3,5)$P_2$. Data are presented as mean ± SEM, *$P < 0.05$ (unpaired t-test).

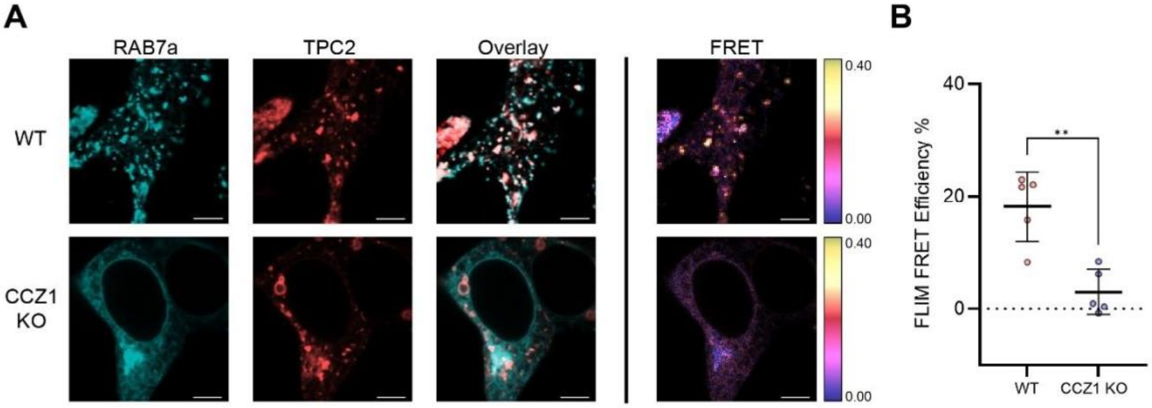


**Supplementary Figure S6: In the absence of apilimod pretreatment, RAB7a–TPC2 interaction is retained in WT but abolished in CCZ1 KO cells**. (A) HEK293 WT (top row) and CCZ1 KO cells coexpressing mTq2-RAB7a (cyan) and TPC2-mVenus (red), with corresponding FLIM-FRET efficiency maps shown on the right. FRET efficiencies (0–40%) are color-coded according to the color scale. Scale bars: 5 µm. (B) FRET efficiencies measured in representative regions of interest (FRET-positive vesicles in WT cells and representative RAB7a-positive areas in KO cells) corresponding to images in A (n=5 WT, n=5 CCZ1 KO). Error bars: ±SD. Statistical significance: Unpaired two tailed t-test, **P < 0.01.

### Fig. S7

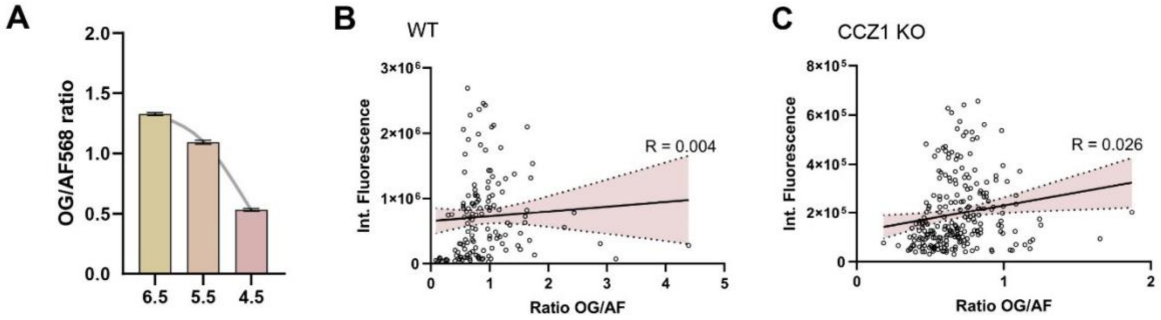


**Supplementary Figure S7: Fluorometric determination of intravesicular pH.** (**A**) The fluorescence intensity ratio per endosome for OregonGreen-Dextran (OG) and Alexa Fluor568-Dextran (AF568) was measured 10 min. after pH clamping cells with Nigericin and Monensin. pH 7.5 was measured as well and showed no difference to pH 6.5. (**B, C**) Integrated Alexa Fluor 568 intensity per endosome was plotted against the OG/AF568 ratio to assess whether dextran uptake influences the ratiometric pH readout. No meaningful correlation was detected (R values shown)

### Fig. S8

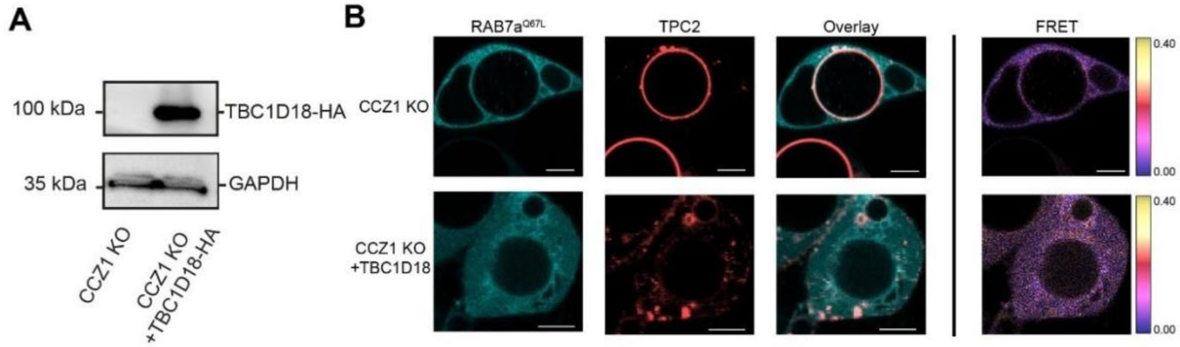


**Supplementary Figure S8: Overexpression of TBC1D18 cannot rescue interaction between TPC2 and RAB7a$^{Q67L}$.** (**A**) TBC1D18 was overexpressed in CCZ1 KO cells and its expression confirmed by western Blot. (**B**) CCZ1 KO cells were co-expressing mTq2-RABa (cyan) and TPC2-mVenus (red), with and without (top row) additional transfection of TBC1D18-HA. The cells were treated overnight with apilimod (1 µM). Corresponding FLIM-FRET efficiency maps are shown on the right. FRET efficiencies (0–40%) are colorcoded according to the color scale. Scale bars: 5 µm.

**Fig. S9**

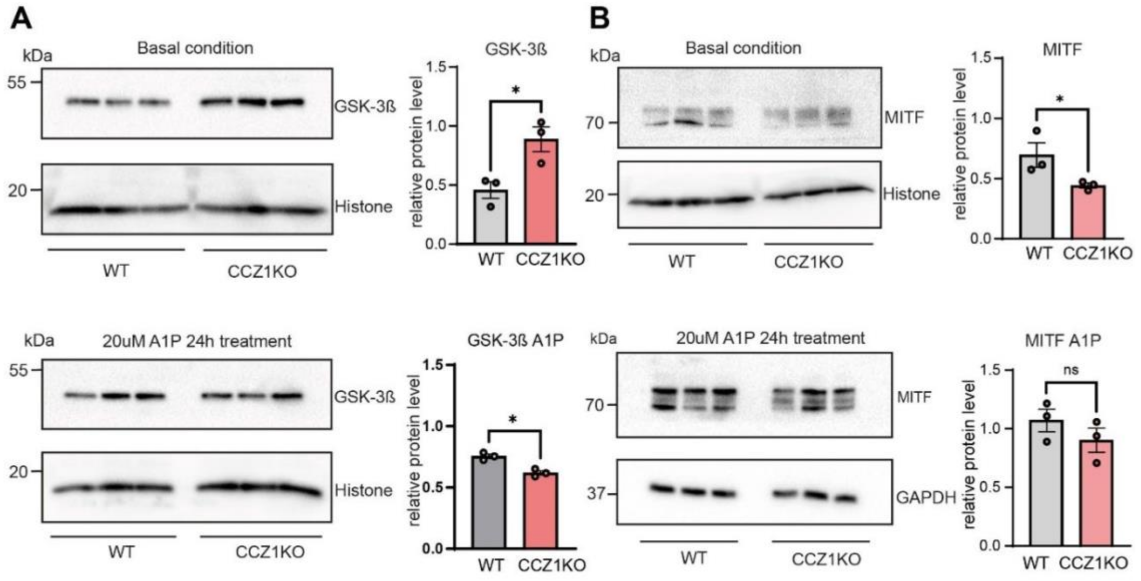


**Supplementary Figure S9: Western blot of GKS-3ß and MITF in SK-MEL5 WT and CCZ1 KO cells under basal and A1P treatment conditions.** (**A**) Western blot of GSK-3ß and MITF under basal condition with statistical analysis on the right side. (**B**) Western blot of GSK-3ß and MITF under after 24h treatment of A1P with statistical analysis on the right side. Band intensity of GSK-3ß and MITF is normalized by loading control. Nonparametric Mann-Whitney test was performed. Each dot represents a biological replicate, median ± indicated interquartile range (IQR) is indicated, *$p < 0.05$, ns: not significant.

**Fig. S10**

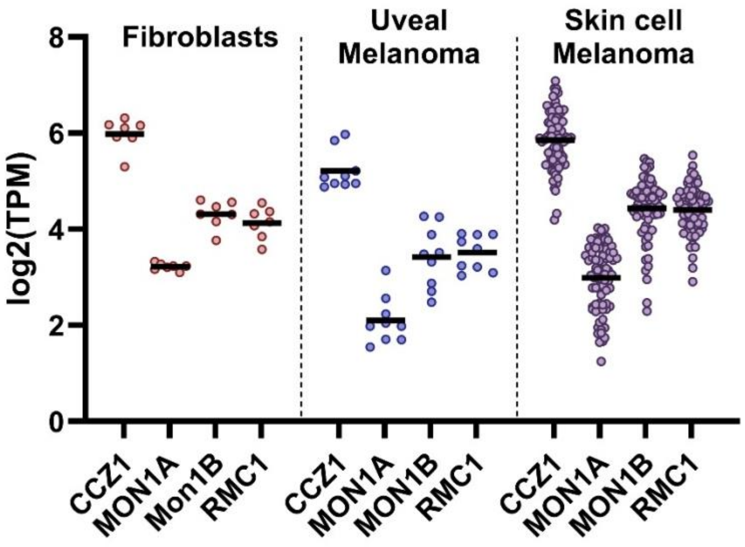


**Supplementary Figure S10: Transcript levels of the CCZ1-MON1a/b-RMC1 complex across fibroblasts and malignant melanoma cell lines.** Steady state RNA levels (log2(TPM+1) (TPM=transcripts per million) of each monomer (CCZ1B, MON1A, MON1B and RMC1) is plotted for healthy fibroblasts, skin cell melanoma, and uveal melanoma cells. Data provided by DepMap (depmap.org).
